# RSV F evolution escapes some monoclonal antibodies but does not strongly erode neutralization by human polyclonal sera

**DOI:** 10.1101/2025.03.11.642476

**Authors:** Cassandra A.L. Simonich, Teagan E. McMahon, Xiaohui Ju, Timothy C. Yu, Natalie Brunette, Terry Stevens-Ayers, Michael J. Boeckh, Neil P. King, Alexander L. Greninger, Jesse D. Bloom

**Affiliations:** Basic Sciences and Computational Biology Divisions, Fred Hutchinson Cancer Center, Seattle, WA 98109; Department of Pediatrics, University of Washington, Seattle, WA, 98195; Pediatric Infectious Diseases Division, Seattle Children’s Hospital, Seattle, WA 98105; Molecular and Cellular Biology Graduate Program, University of Washington and Fred Hutch Cancer Center, Seattle, WA 98109, USA; Department of Biochemistry, University of Washington, Seattle, WA 98195; Institute for Protein Design, University of Washington, Seattle, WA 98195; Department of Laboratory Medicine and Pathology, University of Washington Medical Center, Seattle, WA 98195; Vaccine and Infectious Disease Division, Fred Hutchinson Cancer Center, Seattle, WA 98109; Howard Hughes Medical Institute, Seattle, WA 98109

## Abstract

Vaccines and monoclonal antibodies targeting the respiratory syncytial virus (RSV) fusion protein (F) have recently begun to be widely used to protect infants and high-risk adults. Some other viral proteins evolve to erode polyclonal antibody neutralization and escape individual monoclonal antibodies. However, little is known about how RSV F evolution affects antibodies. Here we develop an experimental system for measuring neutralization titers against RSV F using pseudotyped lentiviral particles. This system is easily adaptable to evaluate neutralization of relevant clinical strains. We apply this system to demonstrate that natural evolution of RSV F leads to escape from some monoclonal antibodies, but at most modestly affects neutralization by polyclonal serum antibodies. Overall, our work sheds light on RSV antigenic evolution and describes a tool to measure the ability of antibodies and sera to neutralize contemporary RSV strains.

## Introduction

Respiratory syncytial virus (RSV) is the leading cause of hospitalization for infants in the United States^1,2^ and can also cause severe illness and death in older adults and immunocompromised individuals^3–5^. RSV has two surface glycoproteins, the fusion (F) and attachment (G) proteins. Neutralizing antibodies targeting the pre-fusion conformation of F are correlated with protection in humans^6–17^. Therefore, recent efforts have focused on developing monoclonal antibodies or vaccines targeting F that can protect infants and high-risk adults. Three vaccines consisting of pre-fusion stabilized F have recently been approved for adults in the USA^10–12,18^, including one to protect infants via passive transfer of maternal antibodies^17,19^. Furthermore, the anti-F monoclonal antibody nirsevimab is now recommended for administration to infants in the USA during RSV season^20–22^. In its first year of use, nirsevimab showed good effectiveness in preventing hospitalizations of infants^13,16,20^, and demand for the antibody outstripped supply^23^. Additional anti-F antibodies are now being developed for use in infants^9,24^. RSV is therefore the first virus for which a monoclonal antibody will be in widespread sustained use in an appreciable fraction of the human population.

Some other human RNA viruses evolve to escape monoclonal antibodies and erode neutralization by polyclonal serum antibodies. For instance, vaccines for influenza virus and SARS-CoV-2 are updated annually to keep pace with viral evolution^25,26^, and over the last few years SARS-CoV-2 escaped most clinically approved monoclonal antibodies^27,28^. The extent to which RSV F evolves to erode neutralization by either polyclonal or monoclonal antibodies is not yet clearly understood. At the sequence level, RSV F is less variable than the surface proteins of influenza virus or human coronaviruses^29^, which has led to less attention to the possibility of antigenic evolution. However, perhaps due to this lack of attention, one monoclonal antibody (suptavumab) failed in an expensive phase III clinical trial because it was escaped by mutations in circulating RSV-B strains^30^. Additionally, recent studies have identified occasional strains with resistance mutations in F to nirsevimab^31–37^.

Efforts to understand how RSV F’s evolution impacts antibodies have been hampered by the lack of assays to easily measure neutralization of F proteins from recent clinical strains. Currently, most common RSV neutralization assays utilize lab-adapted older strains that are not representative of recently circulating viruses^38,39^. Assays utilizing recent clinical strains are challenging because RSV is extremely difficult to grow in the lab^40–45^. Here we overcome these challenges by developing a pseudotyping system to efficiently measure neutralizing antibody titers to the F proteins of historical and contemporary RSV strains. We show that RSV’s natural evolution only modestly affects neutralization by polyclonal serum antibodies, but does lead to escape from some monoclonal antibodies.

## Results

### Efficient pseudotyping of lentiviral particles with F and G from a variety of RSV strains

A widely used approach to study antibody neutralization targeting viral proteins involved in cell entry is to generate single-cycle infectious lentiviral particles that display the viral proteins on their surface and rely on the function of these proteins to infect target cells^46,47^. These “pseudotyped” viral particles can be used to study the effects of genetic variation on the function and antigenicity of the viral proteins at biosafety level 2. A few papers have reported pseudotyping RSV F and G on lentiviral particles^48–51^, but those methods have not been widely used outside the papers themselves, and we were unable to generate appreciable infectious titers of F and G pseudotyped lentiviral particles by following the methods described in those studies (data not shown).

We initially focused on a lab-adapted, subtype A strain of RSV termed the “Long strain”. To pseudotype RSV F and G on lentiviral particles, we transiently transfected 293T cells with expression constructs encoding codon-optimized F and G from the Long strain, lentivirus helper plasmids, and a lentivirus backbone plasmid encoding luciferase and ZsGreen^52^ (Figure 1A). Prior studies of paramyxoviruses report improved pseudovirus titers with truncation of the cytoplasmic tail of the viral proteins^53–55^. For RSV, we first evaluated tiled truncations of G’s N-terminal cytoplasmic tail, which were each paired with full-length F. Titers measured on 293T target cells using a luciferase-based readout increased with longer truncations of the G cytoplasmic tail, and highest titers were achieved with a 31-amino-acid deletion (Figure 1B). We also tiled truncations of F’s C-terminal cytoplasmic tail and paired these with the optimal G (31-residue cytoplasmic tail deletion). Titers were highest for conditions with the full-length F and shorter cytoplasmic tail truncations, but decreased with deletion of 16 or more amino acids of the F cytoplasmic tail (Figure 1C). Therefore, for all subsequent experiments in this paper, we used full-length F and G with a 31-amino-acid cytoplasmic tail deletion. Consistent with prior studies of RSV, we found that flash freezing on dry ice and thawing at 37°C maintains pseudovirus infectivity (Supplemental Figure 1)^49,56^.

**Figure 1.**
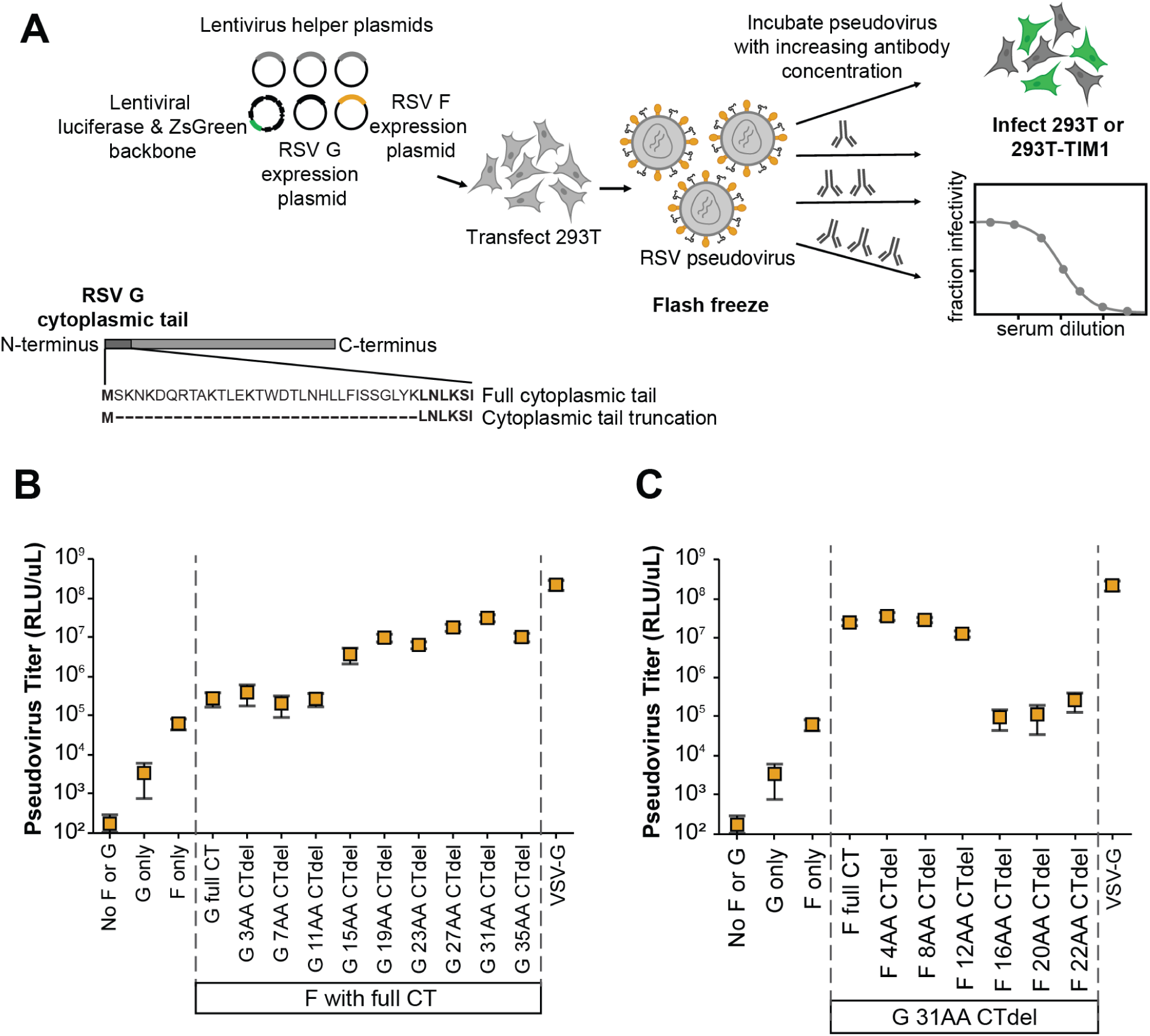
Generation of pseudoviruses with RSV F and G. (A) Process to create lentiviral particles pseudotyped with RSV F and G and then perform neutralization assays. To produce the pseudotyped particles, 293T cells are transfected with lentivirus helper plasmids (encoding Gag-Pol, Tat, and Rev) along with a lentiviral backbone plasmid (encoding luciferase and ZsGreen), and plasmids expressing codon-optimized F and G with a truncated N-terminal cytoplasmic tail (CT). Note G is a type II membrane protein, so the cytoplasmic tail is at the N-terminus. (B) Titers of pseudotyped lentiviral particles expressing the lab-adapted subtype A Long strain F and G increased when full-length F was paired with G with a N-terminal cytoplasmic tail truncation. The highest titers were achieved with a 31-amino-acid truncation of the tail; note all subsequent experiments in this paper use G with this truncation. The plot shows particles pseudotyped with VSV-G as a positive control. Titers are reported in relative luciferase units per microliter (RLU/uL) following infection of 293T cells. Points indicate the mean ± standard error of two independent replicates. (C) Titers of pseudotyped lentiviral particles decreased with C-terminal cytoplasmic tail truncations of the F paired with the truncated G. Points indicate the mean ± standard error of two independent replicates.

We next tested if we could produce high titer pseudoviruses with F and G from other RSV strains in addition to the lab-adapted Long strain. We made pseudoviruses expressing codon-optimized F and G (with the 31-amino acid cytoplasmic tail deletion) from the subtype A lab-adapted A2 and subtype B lab-adapted B1 strains (Supplemental Table 1). The pseudoviruses with F and G from these other two lab-adapted strains also yielded good titers of ∼10^5^ transducing units per milliliter on 293T cells (Figure 2A).

**Figure 2.**
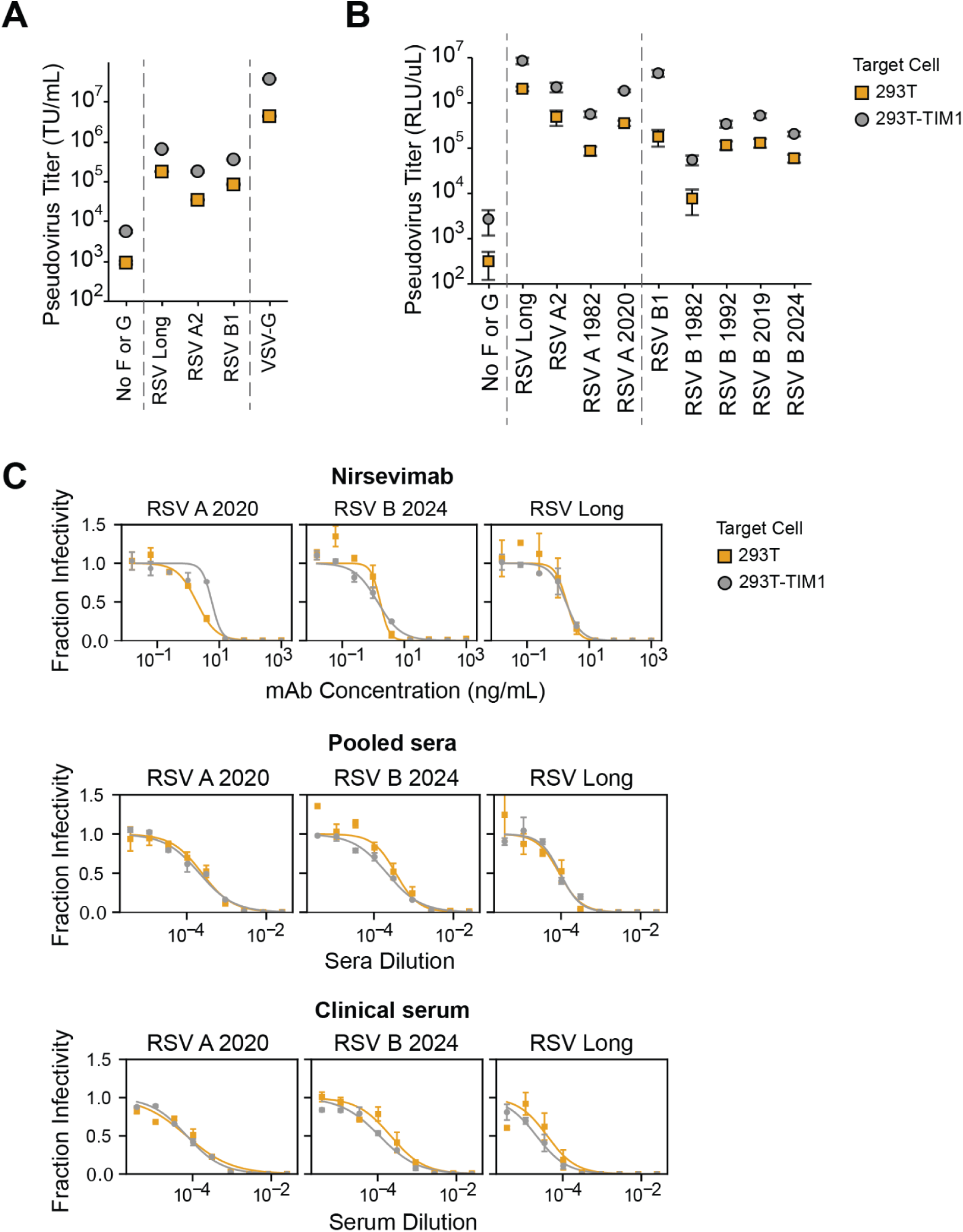
TIM1 expression increases infection of 293T cells by RSV pseudovirus without impacting neutralization. (A) Titers of RSV pseudoviruses are higher on 293T cells expressing TIM1 than unmodified 293T cells. Titers are shown for RSV pseudovirus expressing F and G from lab-adapted strains in transducing units per milliliter (TU/mL) using 293T or 293T-TIM1 target cells. VSV-G pseudotyped particles are shown as a positive control. Points indicate the mean ± standard error of two independent replicates. (B) F and G from a variety of subtype A and B lab-adapted and clinical strains are efficiently pseudotyped on lentiviral particles. Titers in relative luciferase units per microliter (RLU/uL) measured on 293T or 293T-TIM1 cells for RSV pseudoviruses expressing F and G from lab-adapted strains (subtype A Long and A2, subtype B B1), or F from clinical sequences (A 1982, A 2020, B 1982, B 1992, B 2019, B2024) paired with Long G. Points indicate the mean ± standard error of two independent replicates. (C) Neutralization titers for RSV pseudoviruses are the same on 293T and 293T-TIM1 target cells. Neutralization curves for RSV pseudoviruses expressing F from clinical strains RSV A 2020 or B 2024 or from the lab-adapted Long strain paired with Long G against the F-directed monoclonal antibody Nirsevimab, recent pooled human sera, and a human serum specimen collected from a healthy adult in 1987. Points indicate the mean ± standard error of technical replicates.

Because F is the main target of monoclonal antibodies and vaccines^6^, we next tested if we could produce pseudoviruses with F from clinical rather than lab-adapted RSV strains. We chose F proteins from direct sequencing of clinical specimens including subtype A strains from 1982 and 2020 and subtype B strains from 1982, 1992, 2019 and 2024 (Supplemental Table 1). We paired the F proteins (expressed from codon-optimized gene sequences) from clinical strains with the truncated G from the lab-adapted subtype A Long strain, and named the pseudoviruses based on the F protein (RSV A 1982, RSV A 2020, RSV B 1982, RSV B 1992, RSV B 2019 and RSV B 2024; Supplemental Table 1). All of these pseudoviruses with clinical strain F proteins gave appreciable titers on 293T cells (Figure 2B), although the titers were generally lower than for pseudoviruses with F from lab-adapted strains. Pseudoviruses expressing subtype A clinical strain F proteins generally produced higher titers compared to those with subtype B clinical strain F proteins (Figure 2B).

The titers of the RSV F and G pseudoviruses were further enhanced by infecting 293T target cells engineered to overexpress TIM1 (T-cell immunoglobulin and mucin domain 1, Supplemental Figure 2A), which can serve as a non-specific attachment factor for some viruses by binding to phosphatidylserine in the viral membrane^57^. The pseudovirus titers were generally ∼7-fold higher on 293T-TIM1 cells compared to unmodified 293T cells (Figure 2A-B). Note that inclusion of the G protein is less essential for good pseudovirus titers on 293T-TIM1 than 293T cells (Supplemental Figure 2B), but does still enhance titers on both cell lines.

We used pseudoviruses with F from three different RSV strains (the lab-adapted Long strain, and the clinical strains RSV A 2020 and RSV B 2024) paired with Long G to generate neutralization curves against the clinically relevant F-directed monoclonal antibody nirsevimab, a pool of recently collected human sera, and a serum sample collected from a healthy adult in 1987. All three RSV pseudoviruses were neutralized by the antibody and sera, and the neutralization was the same on 293T or 293T-TIM1 target cells (Figure 2C). These data suggest TIM1 expression does not affect the measured neutralization titers, so for all subsequent neutralization assays in this study we used 293T-TIM1 target cells since they gave higher titers. We also found that the RSV pseudovirus must be diluted at least 4-fold to make the neutralization titers independent of the pseudovirus dilution factor (Supplemental Figure 3), possibly because undiluted transfection supernatant contains components that affect neutralization.

### Pseudovirus neutralization titers match expected values from a full-virus neutralization assay and for reference sera

We validated our neutralization assay with RSV pseudoviruses by comparing neutralization titers for 28 human serum specimens measured using our RSV A2 pseudovirus to recently published titers from an assay using full-length, replicative A2 virus^58^. The titers measured with the pseudoviruses were nearly identical to those measured with the full-length virus (Figure 3A, Supplemental Figure 4).

**Figure 3.**
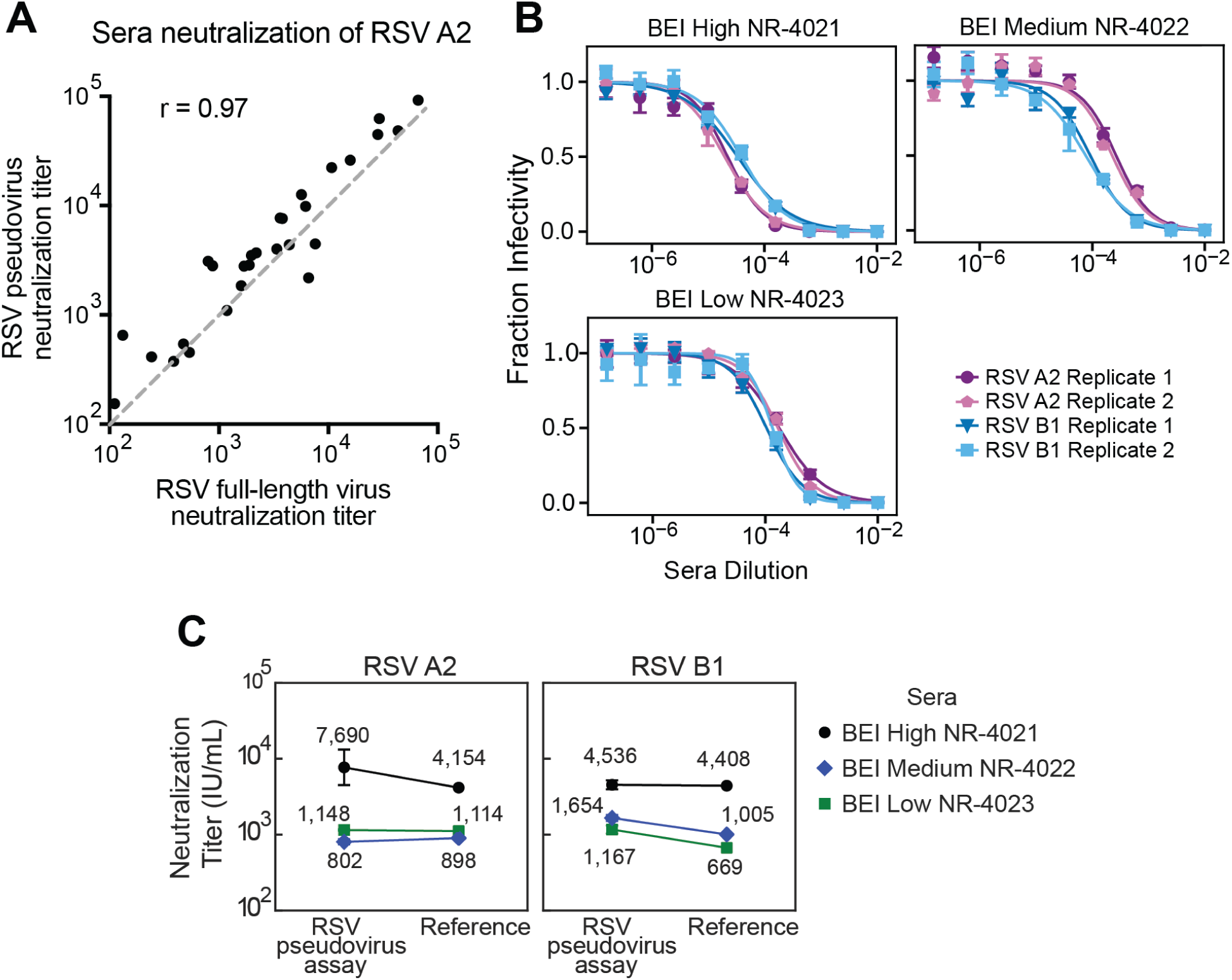
RSV pseudovirus neutralization titers match expected values from a full-virus neutralization assay and reference sera. (A) Correlation between neutralization titers against RSV A2 measured with our pseudovirus neutralization assay (y-axis) versus previously published values measured using full-length replicative RSV A2 (x-axis). 1:1 is shown by the gray dashed line. Neutralization titers are the reciprocal sera dilution at 50% fraction infectivity. Each point represents the geometric mean neutralization titer of two independent experiments for a different human serum sample. (B) Neutralization measurements are highly reproducible. Neutralization curves of pseudoviruses with F and G from the RSV A2 or B1 strains against three different reference sera from BEI Resources. Points indicate the mean ± standard error of technical repeats of experiments performed on the same day. Different replicates indicate independent experiments performed on different days. See Supplemental Figure 5 for additional independent replicates validating the reproducibility. (C) Neutralization titers for the three reference sera from BEI Resources match published values. Plots show neutralization titers for pseudoviruses expressing F and G from RSV A2 or B1 strains versus published neutralization values^58,60^. Neutralization titers were standardized to International Units per milliliter (IU/mL) as described in Supplemental Figure 5.

As further validation, we measured neutralization by three reference sera from BEI Resources against pseudoviruses with F and G from the A2 or B1 strains of RSV. The titers were highly reproducible across independent experiments performed on different days (Figure 3B, Supplemental Figure 5A). To compare neutralization titers of the BEI Resources reference sera to published standardized neutralization titers, we also measured neutralization of the WHO International Standard Antiserum to RSV (NIBSC 16/284) against pseudoviruses with F and G from the A2 or B1 strains of RSV to calculate a conversion factor between pseudovirus neutralization titers and International Units per milliliter (IU/mL)^58–60^ (Supplemental Figure 5A-B). These measurements were also highly reproducible across independent experiments (Supplemental Figure 5A). Standardized RSV pseudovirus neutralization assay titers for the BEI Resources reference sera against both A2 and B1 pseudoviruses were within 2-fold of the published reference values (Figure 3C).

### RSV pseudovirus neutralization assay measures F-directed neutralizing antibodies

Previously published RSV neutralization assays utilizing immortalized cell lines measure neutralization by F-directed but not G-directed antibodies. However, assays on primary cells or using complement on immortalized cell lines can measure G-directed neutralization^39,61–65^. To test the relative contributions of F- and G-directed antibodies to neutralization measured using our pseudovirus assay, we depleted a pool of recently collected human sera of G- or F-directed antibodies (Figure 4A-B) and then measured neutralization of pseudoviruses with F and G from the Long strain (Figure 4C). Depletion of F-directed antibodies dramatically decreased serum neutralization by over two orders of magnitude (Figure 4C). In contrast, depletion of G-directed antibodies caused no appreciable change in neutralization. Therefore, the RSV pseudovirus neutralization assay measures serum neutralization that is almost entirely due to F-directed antibodies.

**Figure 4.**
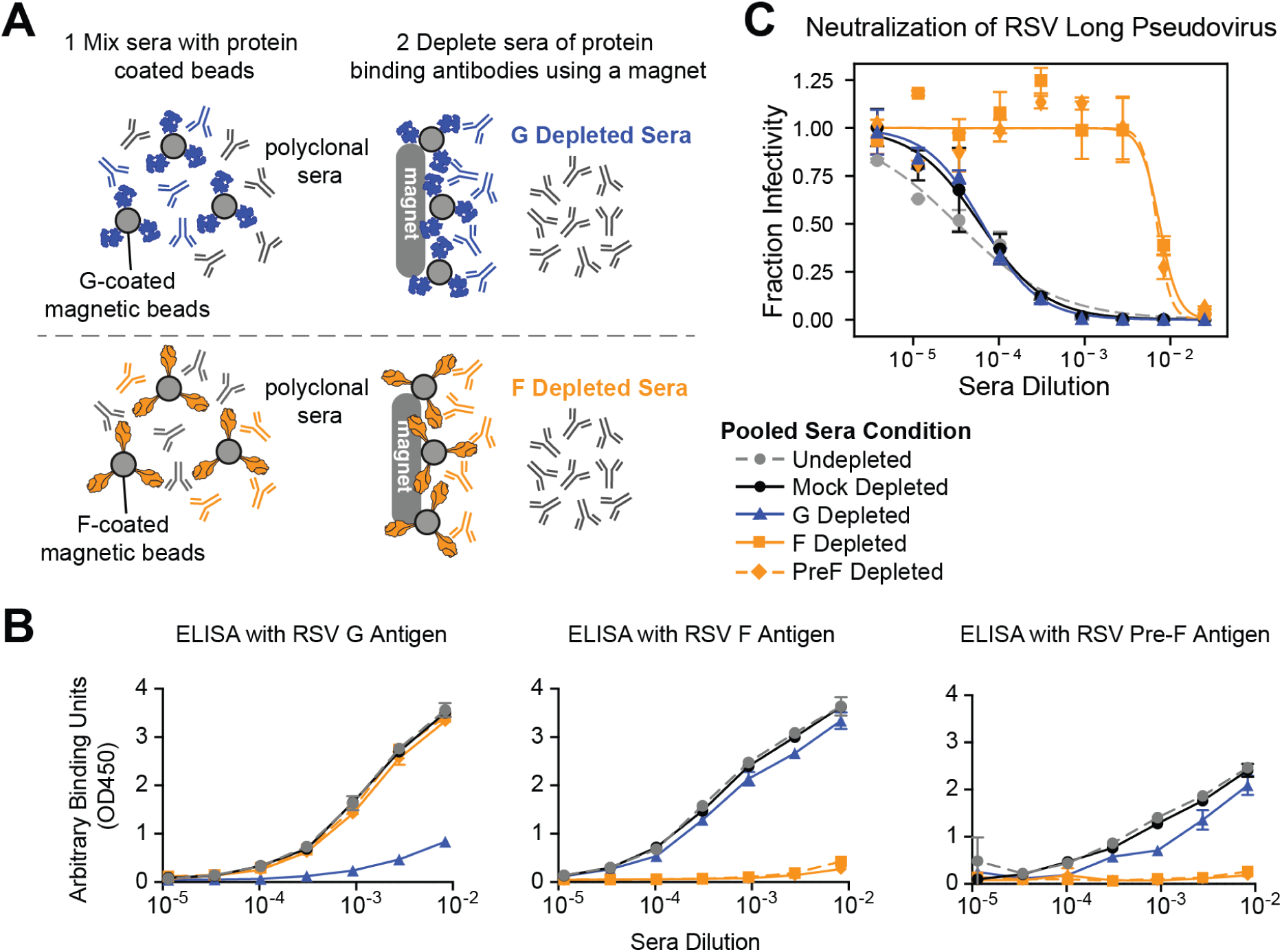
RSV pseudovirus neutralization assay measures F-directed neutralizing activity. (A) Process to deplete pooled polyclonal human sera of either G- or F-binding antibodies using G- or F-coated magnetic beads. Sera was depleted of G-directed antibodies using beads coated with the Long G protein, and depleted of F-directed antibodies using beads coated with either unmodified or pre-fusion stabilized (PreF) Long F protein. (B) ELISA binding curves validate that the depletions nearly completely removed the G- or F- binding antibodies. The ELISA plates were coated with G, unmodified F, or pre-fusion stabilized F all from the Long strain. Points indicate the mean ± standard error of two technical replicates. (C) Neutralization titers against pseudoviruses with the Long strain F and G decrease with depletion of F- but not G-directed antibodies. Shown is the neutralization of pseudovirus with Long F and G against sera depleted by the indicated method, or undepleted or mock depleted sera. Points indicate the mean ± standard error of two technical replicates. All experiments shown in this figure utilized a pool of human sera collected in 2021.

### RSV F evolution does not erode polyclonal sera neutralizing titers

To determine if the evolution of RSV F erodes neutralization by human polyclonal serum antibodies, we measured neutralization of pseudoviruses with F from historical and recent RSV strains. Specifically, we used the F proteins from 1982 and 2020 strains of subtype A, or 1982, 1992, 2019, and 2024 strains of subtype B (Figure 5, Supplemental Table 1). The sequences for these proteins are all close to the trunk of the global RSV F phylogenetic trees rather than on long branches (Figure 5), indicating that they are representative of strains from that time and do not have rare private mutations. The 1982 and 2020 subtype A proteins differ by only two amino-acid mutations, both in defined antigenic regions^66^(Figure 5A). The 1982 and 2024 subtype B proteins differ by ten amino-acid mutations, eight of which are in antigenic regions (Figure 5B).

**Figure 5.**
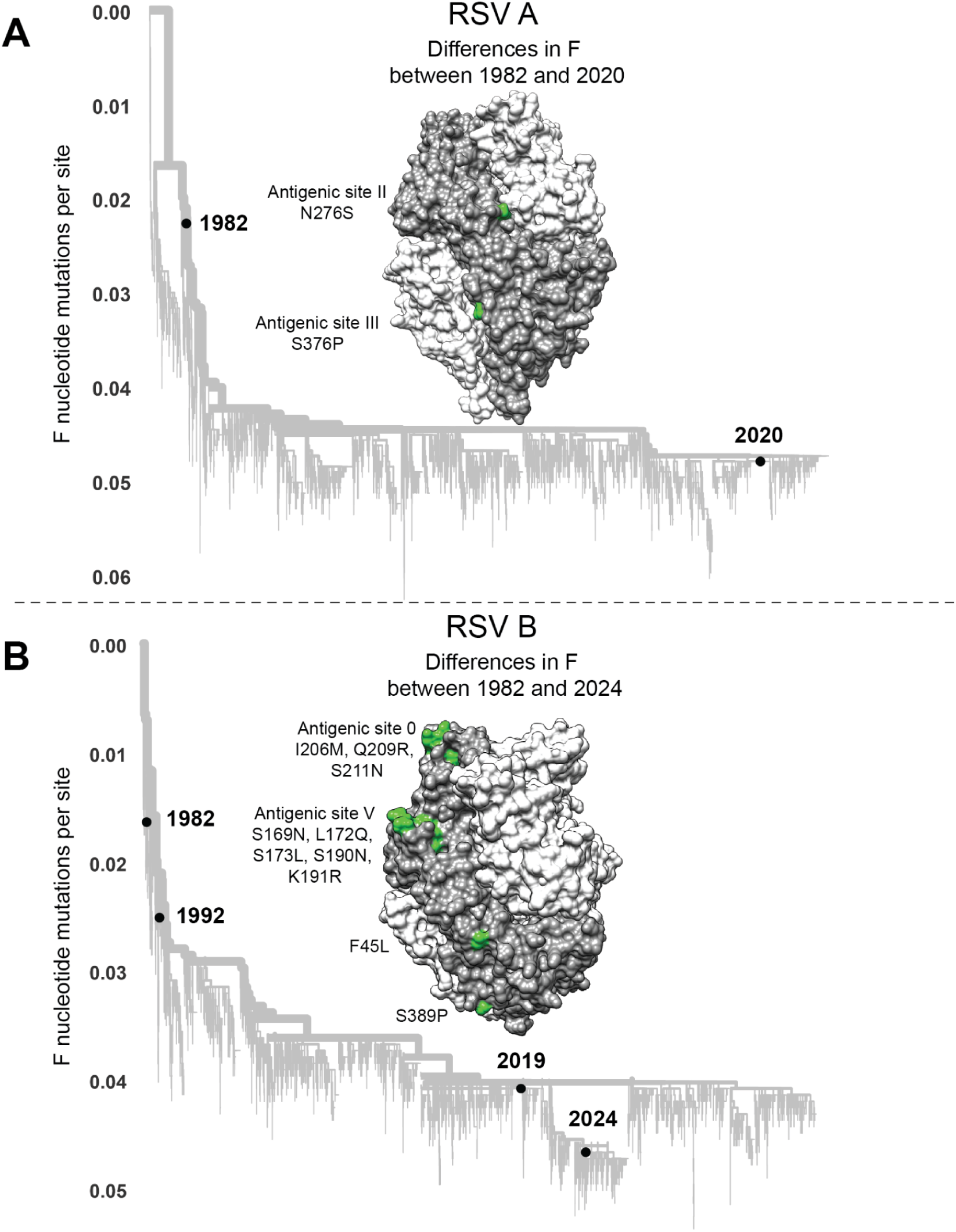
RSV strains used to study effects of F evolution on serum and antibody neutralization. (A) Phylogenetic tree of subtype A RSV F gene sequences with black circles indicating the historical (1982) and recent (2020) sequences selected for study. Amino-acid differences in F between the historical 1982 and recent 2020 sequences are shown in green on one protomer of the RSV F homotrimer (PDB 5C6B); the protomer with the amino-acid differences indicated is in dark gray and the other two protomers are in light gray. An interactive version of this phylogenetic tree can be found here https://nextstrain.org/community/jbloomlab/RSV-evolution-neut@main/RSV-A-F. (B) Phylogenetic tree of subtype B RSV F gene sequences with black circles indicating the historical (1982 and 1992) and recent (2019 and 2024) sequences selected for study. Amino acid differences in F between the 1982 and 2024 sequences are shown on the F structure as in panel (A). An interactive version of this phylogenetic tree can be found here https://nextstrain.org/community/jbloomlab/RSV-evolution-neut@main/RSV-B-F.

To test the effect of F’s evolution on serum antibody neutralization, we used the approach in Figure 6A. We assembled a panel of 20 historical sera collected from healthy adults between 1985 and 1987 (Supplemental Table 2). No information was available about recent respiratory virus infections of these individuals, but since humans are regularly re-infected with RSV throughout adulthood^3,67–69^, most of these individuals were likely infected with a 1980s RSV strain within a few years preceding serum collection. None of the individuals would have been infected with recent strains prior to serum collection, since those strains did not yet exist. We depleted all sera of antibodies directed to G (from the Long strain) to ensure any measured neutralization was due to F-binding antibodies. We then measured neutralization by each historical serum of pseudoviruses with F from the historical and recent strains paired with G from the Long strain. If F evolution erodes serum antibody neutralization then historical sera should neutralize historical F strains better than recent F strains, otherwise the strains should be similarly neutralized (see hypothetical data in Figure 6A).

**Figure 6.**
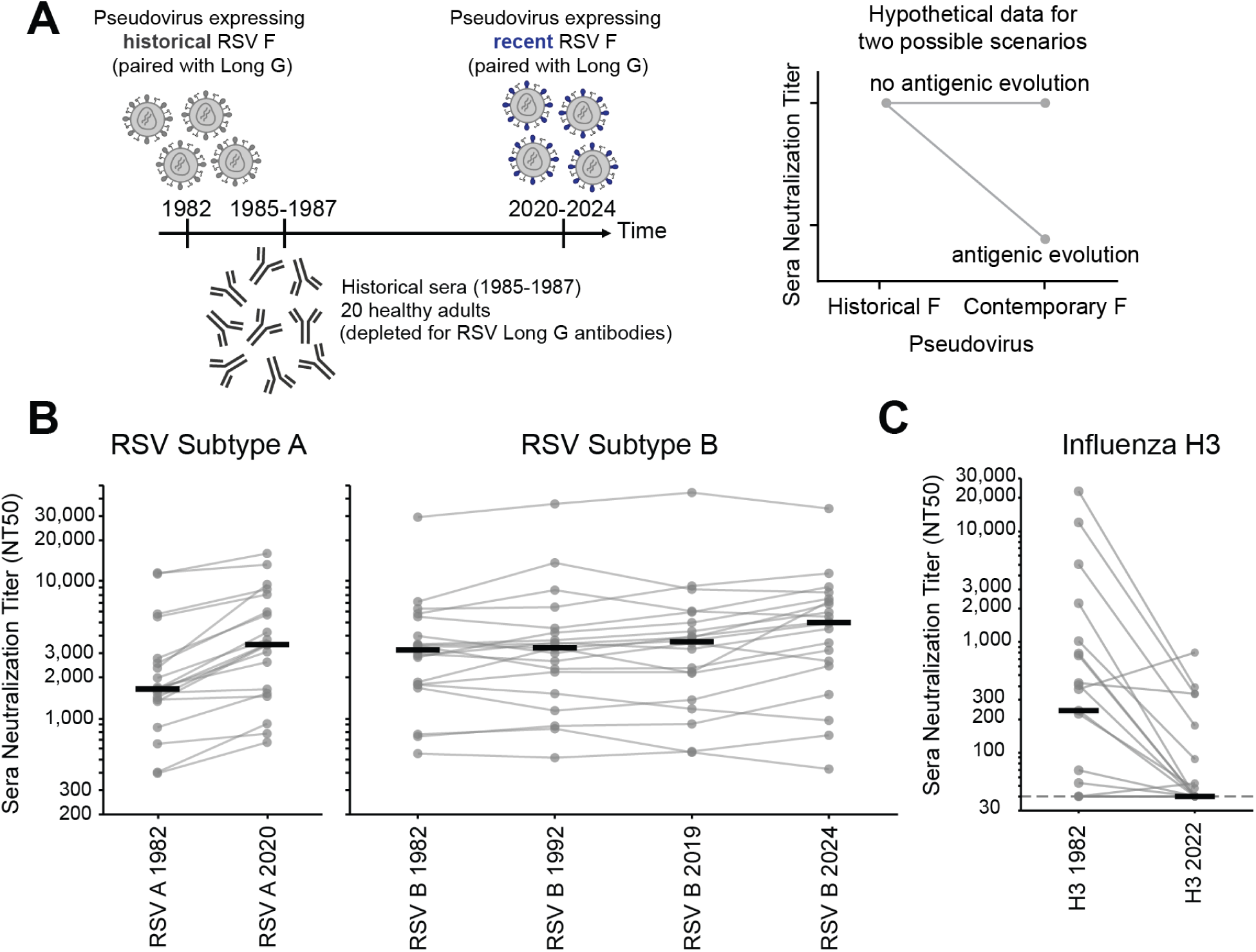
RSV F evolution does not strongly erode polyclonal serum antibody neutralization. (A) Design of experiment to test if F evolution erodes serum antibody neutralization. Historical sera collected from healthy adults in 1985-1987 were depleted for G-binding antibodies, and then tested for neutralization of RSV pseudoviruses with G from the Long strain and F from either an historical (eg, 1982) or a recent (eg, 2020-2024) subtype A or B strain. The plot at right shows hypothetical data for two different scenarios: the sera are expected to neutralize pseudoviruses with the historical F (since it is from individuals who would have been infected with similar viruses), but will not neutralize pseudoviruses with the recent F if that protein undergoes rapid antigenic evolution. (B) Neutralization titers for 20 historical sera against the pseudoviruses expressing the Long strain G and F from historical and recent RSV A (left panel) or RSV B strains (right panel). Each set of gray dots connected by lines represent titers against a different serum (geometric mean of two independent experiments), and the black lines represent the median titer against that pseudovirus across all 20 sera. Individual neutralization curves are in Supplemental Figure 6. (C) Neutralization titers decrease for the same 20 historical sera against pseudoviruses expressing the hemagglutinin from an historical (A/Netherlands/233/1982) versus recent (A/Massachusetts/18/2022) strain of human H3N2 influenza. Dots and lines have the same meaning as in panel (B). Individual neutralization curves are in Supplemental Figure 7.

Neutralization by historical sera was similar for pseudoviruses expressing historical and recent F proteins from both RSV subtype A and B (Figure 6B, Supplemental Figure 6). Specifically, most individual sera neutralized pseudoviruses with F proteins of the 1982 and 2020 subtype A strains at similar titers, and likewise for pseudoviruses with the F proteins of subtype B from 1982, 1992, 2019, and 2024 strains (Figure 6B). This lack of erosion of serum neutralization by RSV F evolution contrasts to similar experiments with the same sera against pseudoviruses with influenza hemagglutinin from historical versus recent human H3N2 strains, which show that influenza hemagglutinin evolution largely escapes serum neutralization over the same timeframe (Figure 6C, Supplemental Figure 7). We note that prior experiments on other sera have also shown rapid erosion of neutralization by the evolution of the spikes of CoV-229E and SARS-CoV-2^70,71^. Overall, these results suggest that RSV F evolution does not rapidly erode polyclonal serum antibody neutralization, in contrast to the evolution of influenza hemagglutinin and spike proteins of SARS-CoV-2 and CoV-229E.

### Natural evolution of RSV F escapes some monoclonal antibodies

Even if RSV F’s evolution does not strongly erode neutralization by polyclonal serum antibodies, it could lead to escape from individual monoclonal antibodies. Indeed, sporadic RSV strains have been reported to carry mutations that escape nirsevimab, which is clinically used for RSV prevention for infants^31–34^. These known escape mutations (K68Q/N, N201S/T) are found in a few sequences scattered across the global RSV-B F phylogenetic tree (Figure 7A). We chose F proteins from recent RSV B strains with the K68Q, N201S, and N201T nirsevimab escape mutations to test for neutralization by monoclonal antibodies, also testing the more representative historical and recent RSV F proteins described in the previous section.

**Figure 7.**
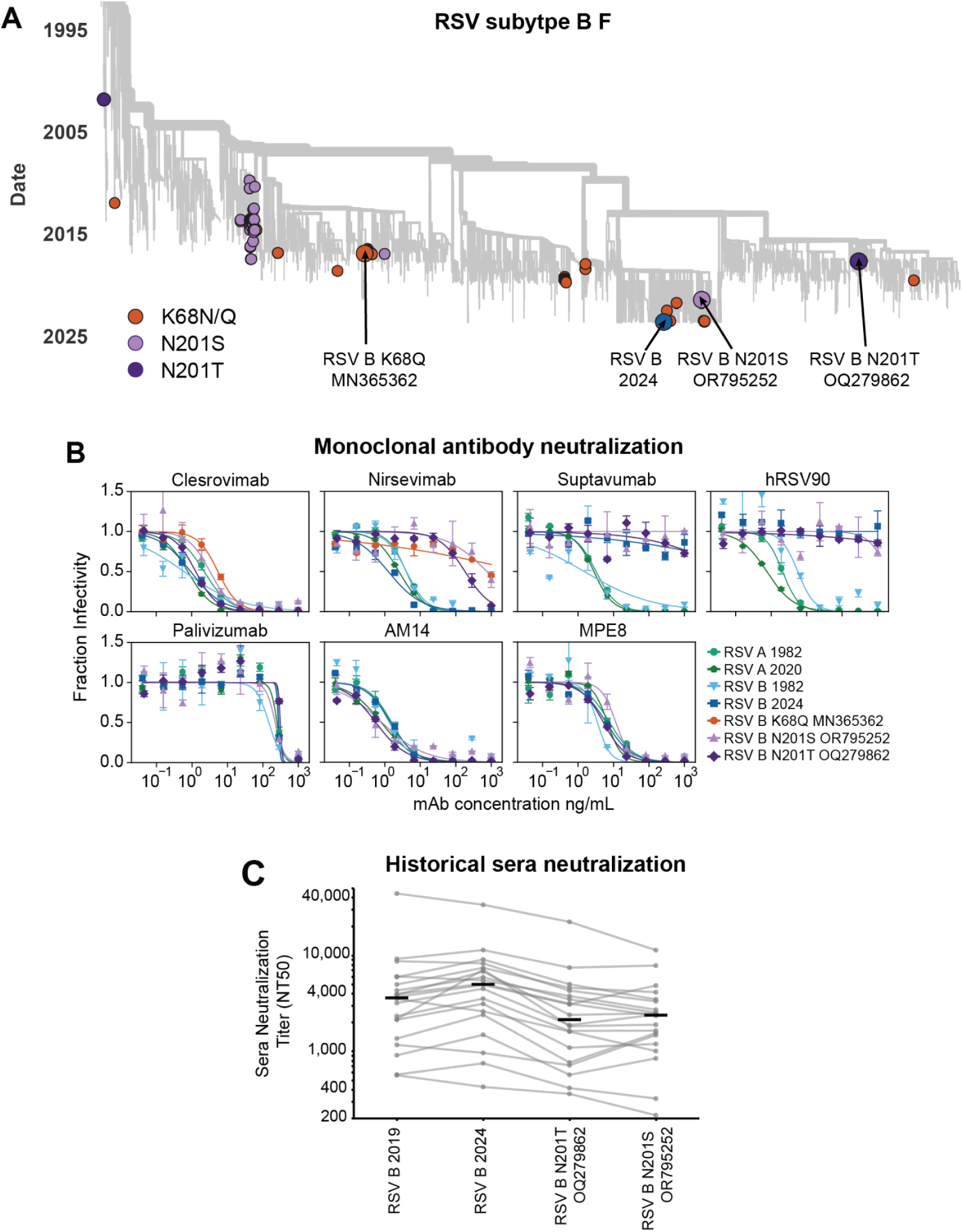
Natural evolution of RSV F escapes some monoclonal antibodies. (A) Phylogenetic tree of F gene sequences from recent subtype B RSV strains, with colored circles indicating the representative RSV B 2024 strain used in our experiments as well as other strains containing known nirsevimab resistance mutations. Arrows indicate F proteins used in our experiments. (B) Neutralization curves for monoclonal antibodies versus pseudoviruses expressing the indicated F protein paired with Long G. Points indicate the mean ± standard error of technical replicates. Neutralization curves for these monoclonal antibodies from a separate experiment performed on a different day as well as versus pseudoviruses expressing F and G from lab-adapted strains are in Supplemental Figure 8. (C) Neutralization titers for the 20 historical sera against pseudoviruses expressing the indicated F protein paired with Long G. Each set of gray dots connected by lines represent titers against a different serum (geometric mean of two independent experiments), and the black lines represent the median titer against that pseudovirus across all 20 sera. The historical sera are the same as those used in Figure 6B,C, and are depleted of antibodies that bind to Long G. The points for the RSV B 2019 and RSV B 2024 strains shown in this panel are the same as those shown in Figure 6B. See Supplemental Figures 6 and 9 for the neutralization curves.

Pseudoviruses expressing the representative F proteins from both recent and historical RSV A and B strains were all neutralized by nirsevimab as well as four other monoclonal antibodies that are in clinical use (palivizumab^72,73^), clinical trials (clesrovimab^74^) or pre-clinical studies (AM14^75^, and MPE8^76^) (Figure 7A, Supplemental Figure 8). However, two other antibodies (suptavumab^30^ and hRSV90^77^) failed to neutralize the representative recent RSV B strain (Figure 7A, Supplemental Figure 8). We note that the failure of suptavumab to neutralize recent RSV B strains was already known, but was discovered only through the costly failure of a phase III clinical trial of this antibody in humans^30^.

Pseudoviruses expressing F from recent RSV B strains with the K68Q, N201S, or N201T mutations were strongly resistant to nirsevimab, and also escaped suptavumab and hRSV90 (Figure 7A, Supplemental Figure 8). These results highlight the potential for recent RSV B F variants to evolve to escape neutralization by a monoclonal antibody in clinical use, and emphasize the importance of using assays like the one we describe here to monitor for resistance in recent strains.

We next evaluated whether the recent F proteins with the sporadic mutations that escape nirsevimab also have reduced neutralization by polyclonal human sera. To do this, we tested neutralization of the pseudoviruses expressing recent RSV B F proteins with the N201S/T mutations against the 20 historical sera described in the prior section (again depleted of G antibodies). A number of the sera showed modestly reduced neutralization of F with the N201S/T mutations relative to the F of representative recent RSV B strains that lack these mutations (Figure 7C; Supplemental Figure 9). Therefore, for some sera a measurable fraction of the neutralization is due to antibodies targeting an epitope similar to that bound by nirsevimab, and so nirsevimab resistance mutations can also influence serum neutralization.

## Discussion

We developed a pseudotyping system to quantify how evolution of RSV F affects antibody neutralization. We found that efficient pseudotyping of RSV F and G on lentiviral particles requires truncation of the G cytoplasmic tail, and that titers were further improved by the use of TIM1-expressing target cells. With this system, we produce high titers of pseudovirus expressing F from clinical strains of both RSV subtype A and B. Neutralization titers measured using these pseudoviruses closely match those from a full-length replicative virus assay.

We found that the evolution of F even over several decades only modestly affects neutralization by human polyclonal serum antibodies. Therefore, RSV F stands in clear contrast to the entry proteins of human seasonal influenza and coronaviruses, which evolve to rapidly erode serum antibody neutralization^25,70,71^. Further work will be needed to determine why RSV F has a slower rate of antigenic change than some other viral entry proteins, but possible hypotheses are that F is less tolerant of mutations^78,79^ or that polyclonal serum antibodies target so many distinct F epitopes that individual mutations are unable to provide much antigenic benefit to the virus^80^. Despite the lack of strong erosion of F-directed neutralization by RSV evolution, humans are reinfected by RSV multiple times throughout their lives^2,3,69,81^. The fact that these reinfections occur despite F’s relatively slow antigenic evolution suggest that either F-directed neutralizing antibody titers wane over time^82–85^, or that F-directed neutralizing antibodies are not actually as protective as widely believed—although this second explanation would seem at odds with studies indicating good efficacy of anti-F vaccines and monoclonal antibodies in humans^6–17^. In either case, the fact that reinfections can occur even without much F antigenic evolution could in turn lead to less immune pressure on F to evolve antigenically.

However, our results confirm that the natural evolution of F does result in escape from some individual monoclonal antibodies. In particular, we confirmed that more recent subtype B strains have escaped monoclonal antibodies (eg, suptavumab and hRSV90) that target a prefusion-specific antigenic region near the trimer apex, consistent with prior studies^30,86^. The fact that monoclonal antibody nirsevimab is now being widely administered to infants in the USA and some other countries could increase the evolutionary pressure on F^20,87^. We confirmed that rare RSV B strains have sporadic mutations that confer escape from nirsevimab^31–34^. Nirsevimab targets an epitope on the apex of the F trimer that is relatively close to the region targeted by the antibodies suptavumab and hRSV90 that are now escaped by most recent subtype B strains^30,77,88^. Therefore, it may be advisable to consider cocktails with additional antibodies targeting other RSV epitopes. We also note that the antibody escape mutations observed to date seem to be mostly in RSV B^30–34^, which has accumulated more mutations than RSV A over the last few decades.

We found that the currently rare nirsevimab escape mutations in some RSV B strains also measurably reduced F-directed neutralization by the polyclonal antibodies in some sera. This finding suggests that an appreciable fraction of neutralization by some sera is due to antibodies targeting the apex of the F trimer near the nirsevimab site^8,66^. Therefore, as nirsevimab (and possibly future monoclonal antibodies) are more widely administered, it will be important to monitor not only for resistance to the antibodies themselves, but also evaluate if any resistance mutations affect neutralization by polyclonal antibodies elicited by infection or vaccination. The pseudovirus assay we describe here will therefore be a valuable tool to continue to monitor how new antibodies and vaccines affect and are affected by RSV evolution.

### Limitations of Study

Our RSV pseudovirus neutralization assay measures F-directed neutralizing antibodies. The G protein is maintained by RSV in nature so is clearly important for viral infection of airway cells during human infections, but is largely dispensable for viral infection of immortalized cell lines^50,51,89,90^. G is particularly dispensable on the 293T-TIM1 cells used in our paper, likely because TIM1 binding to phosphatidylserine on the virion can mediate attachment^57^. Note that most other established RSV neutralization assays also only measure F-directed neutralization^8,61–64,89^. Despite the inability of our assay to measure G-directed neutralization, G is under stronger pressure for adaptive evolution than F during actual RSV evolution^29^, suggesting G-directed antibodies play an important role in human immunity. Our pseudovirus assay also does not measure other immune mechanisms that could be relevant to RSV immunity, including T-cells and antibody effector functions.

Additionally, the pseudotyped lentiviral particles used in our assays likely have a different morphology and density of F and G glycoproteins than authentic RSV, which could influence some aspects of antibody neutralization. We also note that the nucleolin protein thought to act as a co-receptor for F may be more highly expressed in cell lines like the ones used for our assay than in primary human airway cells^91,92^.

Our experiments primarily used sera collected in the 1980s from healthy adults, but no information is available about recent respiratory virus infections of these individuals. It is possible that the titers and specificities of serum antibodies could differ in individuals of other ages or exposure histories.

## Acknowledgements

We thank Richard Neher and Laura Urbanska for maintaining the public RSV Nextstrain build used for the phylogenetic trees in this paper. We thank Caleb Carr for assistance with the phylogenetic trees. The human sera from the 1980s is from the Infectious Disease Sciences Biospecimen Repository at the Vaccine and Infectious Disease Division (VIDD) of the Fred Hutchinson Cancer Center. This work was funded by the NIH / NIAID under R01AI141707 (to JDB) and 1U19AI181767 (subcontract to JDB) and by the Gates Foundation under award INV-072143 (to JDB). JDB is an Investigator of the Howard Hughes Medical Institute. CALS is a Fellow in the Pediatric Scientist Development Program. This project was supported by Award Number K12-HD000850 from the *Eunice Kennedy Shriver* National Institute of Child Health and Human Development. TCY was supported by the NSF graduate research fellowship DGE-2140004. This research was supported by the Flow Cytometry Shared Resource, RRID:SCR_022613, of the Fred Hutch/University of Washington/Seattle Children’s Cancer Consortium (P30 CA015704).

## Author Contributions

Conceptualization: CALS, TEM, JDB; Methodology: CALS, TEM, JDB, XJ, TCY, NB; Investigation: CALS, TEM; Resources: XJ, TCY, NB, TSA, MJB, NPK, ALG; Data curation: CALS, TEM, JDB; Visualization: CALS, TEM, JDB; Writing-Original Draft: CALS, TEM, JDB; Writing-Review & Editing: CALS, TEM, XJ, TCY, NB, TSA, MJB, NPK, ALG, JDB; Supervision: JDB; Funding Acquisition: CALS, JDB.

## Supplemental Figures

**Supplemental Figure 1.**
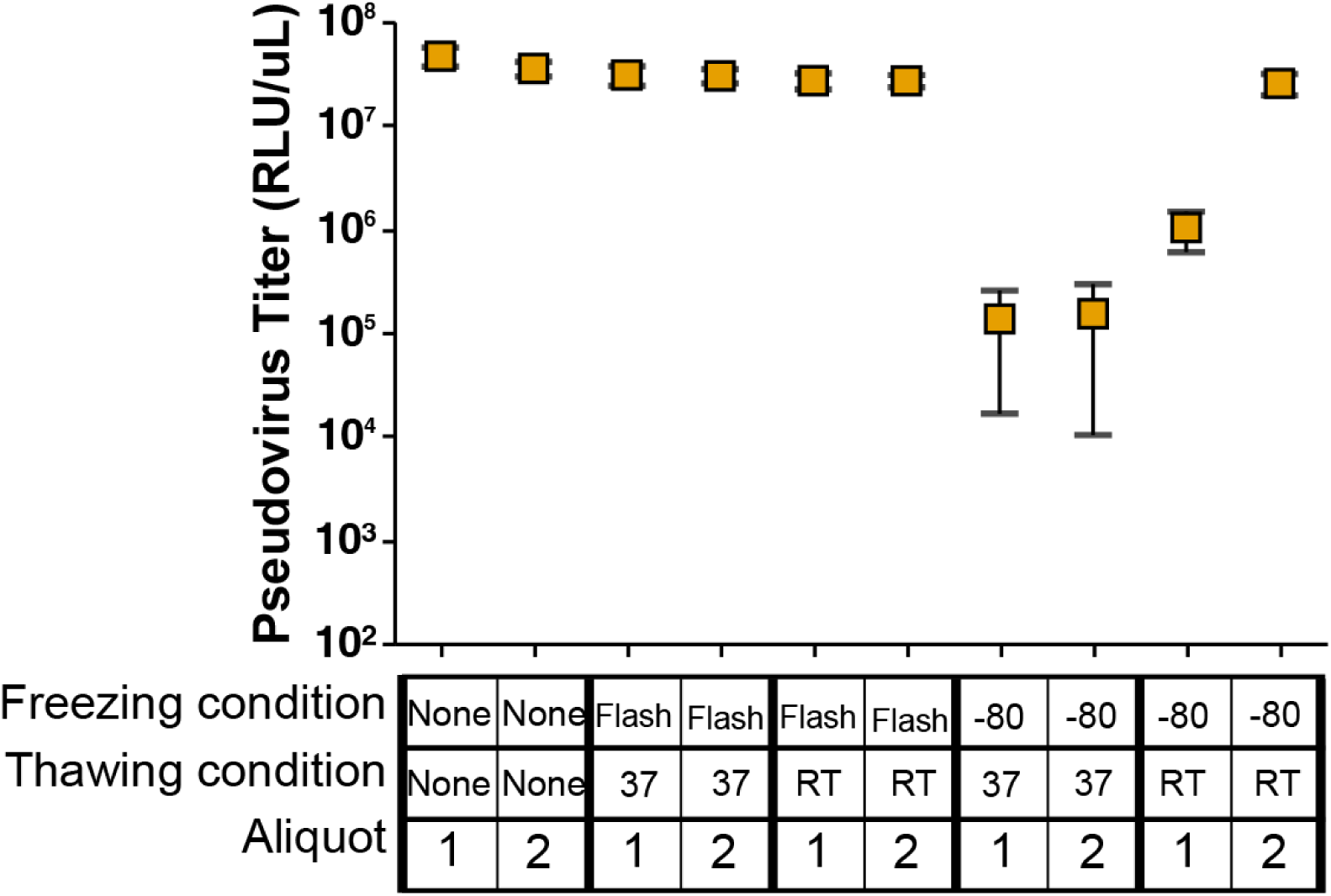
Flash freezing on dry ice maintains pseudovirus infectivity. RSV Long pseudovirus titers (RLU/uL) after different freezing and thawing conditions. Points represent the average titer of two technical replicates. Flash freezing refers to freezing aliquots on dry ice before moving to -80°C storage. -80 freezing refers to placing aliquots directly in the -80°C freezer for a slow freeze. Samples in the 37 thawing condition were thawed in a 37°C water bath while RT samples were thawed at room temperature.

**Supplemental Figure 2.**
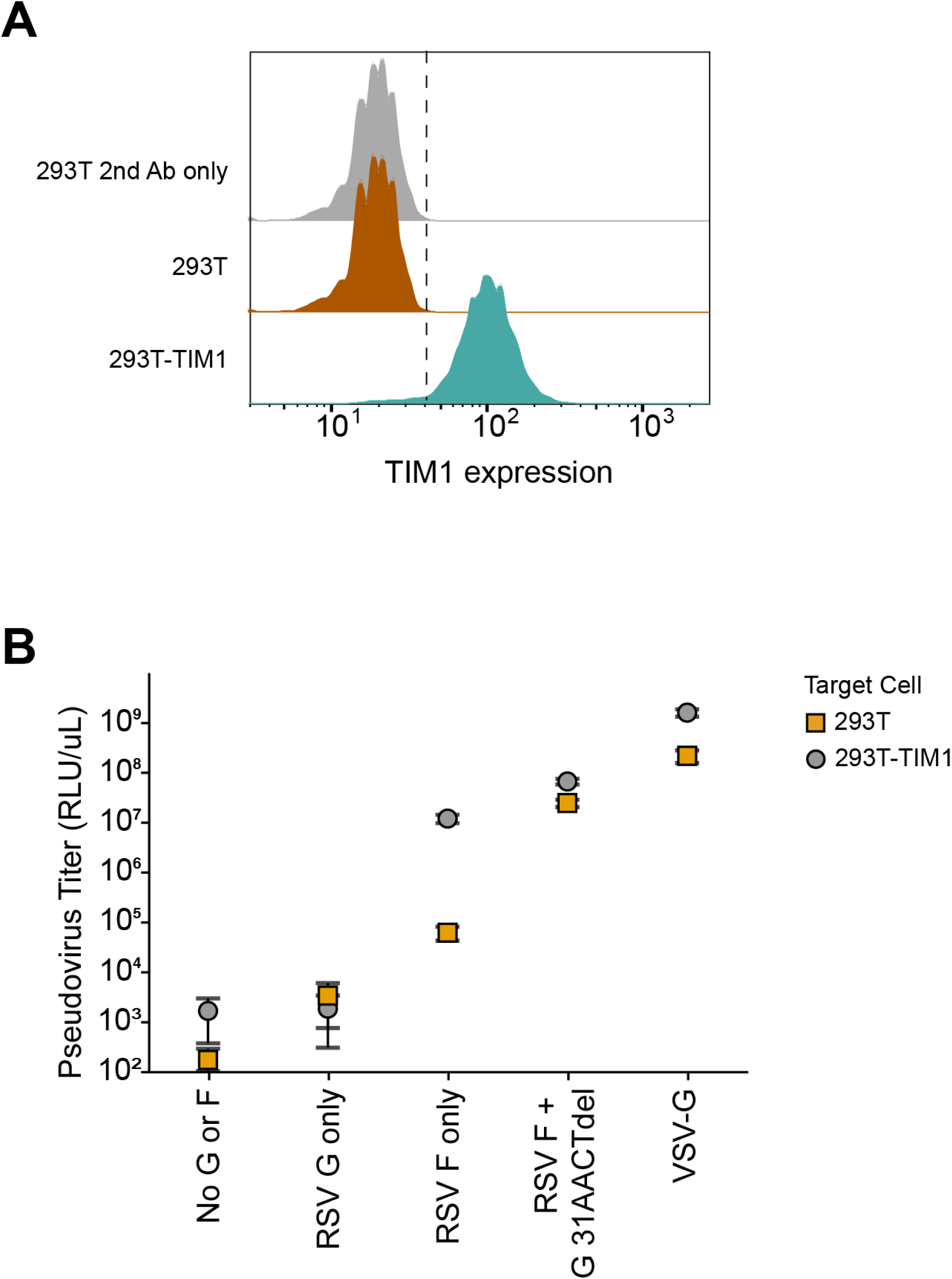
RSV G improves pseudovirus titers even on cells expressing TIM1. (A) Immunostaining confirms the expression of TIM1 in 293T-TIM1 cells. Flow cytometry analysis of 293T cells stained with fluorophore-conjugated secondary antibody only (gray), 293T cells stained with anti-TIM1 primary antibody and secondary antibody (orange) and 293T-TIM1 cells stained with anti-TIM1 primary antibody and secondary antibody (teal). (B) Titers of pseudoviruses expressing the indicated F or G proteins from the Long strain on 293T or 293T-TIM1 target cells. Prior RSV pseudotyping studies have described expression of F alone as being sufficient to produce infectious pseudovirus particles^49–51^. We found that pseudoviruses expressing F alone do infect 293T cells with only low titers, and the addition of the G with a cytoplasmic tail truncation improves titers by 400-fold as shown in Figure 2. This figure shows that infections on 293T-TIM1 cells with pseudovirus expressing F alone results in higher titers, but the addition of the truncated G still improves titers by an additional 6-fold.

**Supplemental Figure 3.**
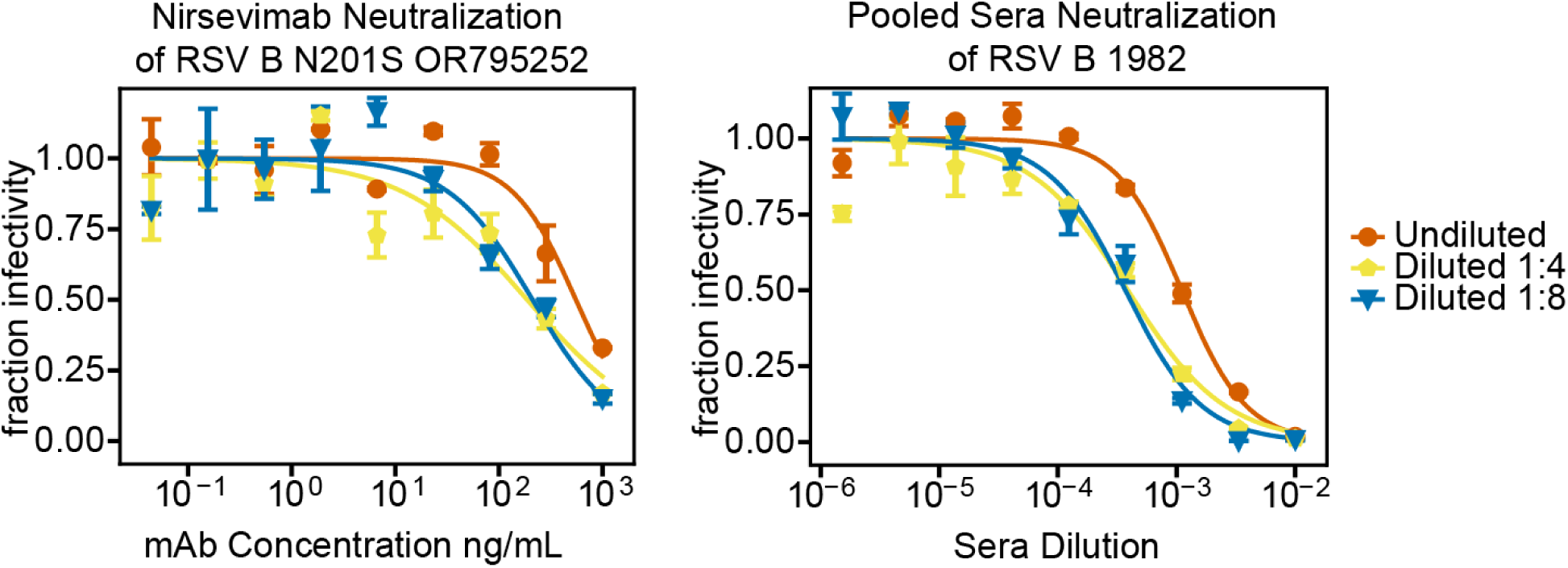
Pseudovirus dilution affects neutralization curves. Neutralization curves using undiluted pseudovirus generated from transfection differ from the curves generated using pseudovirus diluted at least 4-fold for the neutralization assay. Neutralization curves for the monoclonal antibody Nirsevimab versus pseudovirus expressing a clinical RSV B F protein (RSV B N201S OR795252) paired with the Long strain G and pooled human sera collected in 2021 versus pseudovirus expressing a clinical RSV B F protein from 1982 paired with the Long strain G. Different colored curves represent pseudovirus supernatant from the transfected producing cells that was undiluted (orange) or diluted in fresh media (1:4 yellow and 1:8 blue) for the neutralization assay. Points indicate the mean ± standard error of two technical replicates. Based on these findings, for all experiments in the paper we used pseudovirus that had been diluted at least 1:4 after its generation by transfection.

**Supplemental Figure 4.**
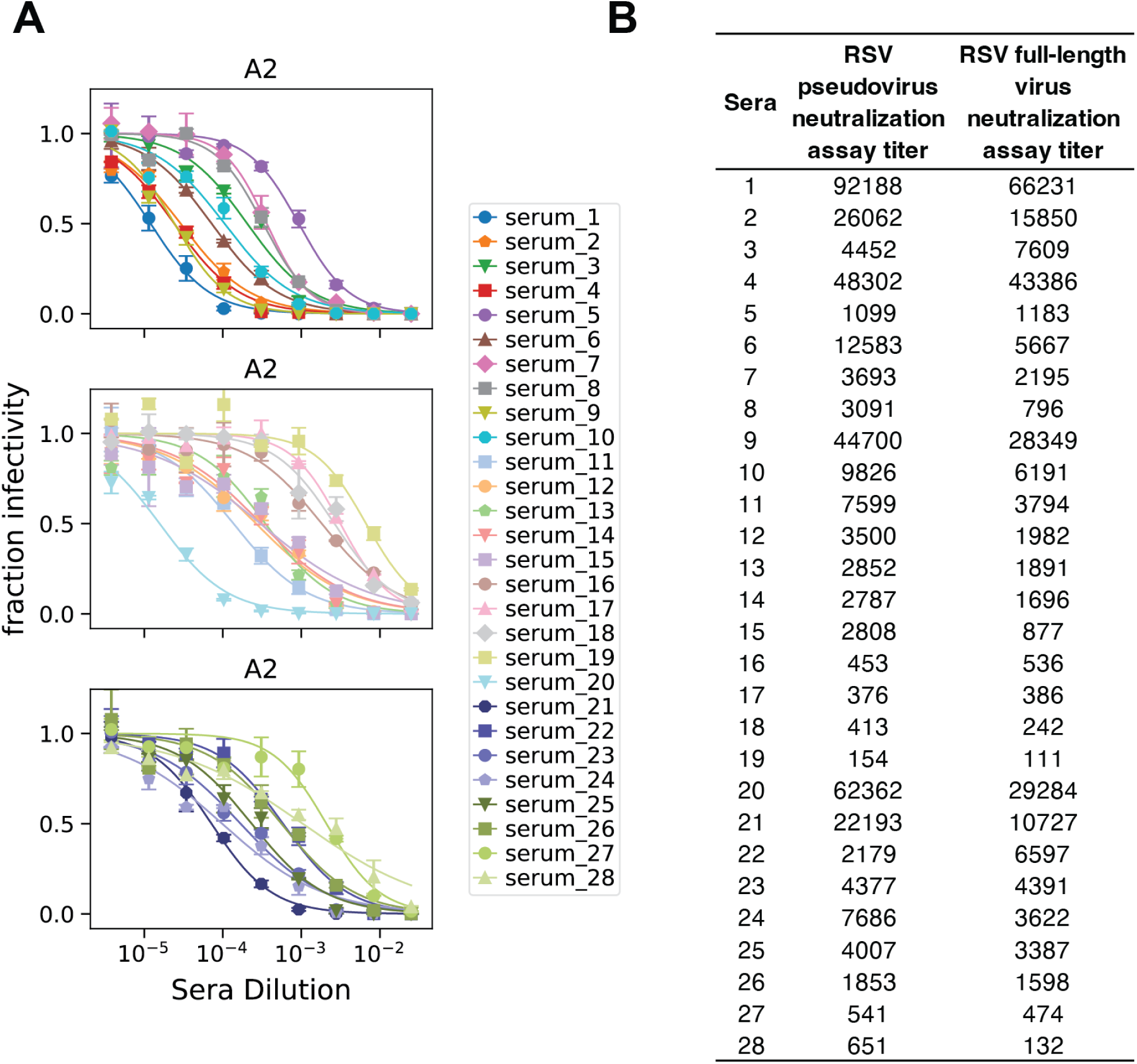
Comparison of RSV A2 pseudovirus and full-length A2 virus neutralization. (A) Neutralization curves for RSV A2 pseudovirus versus 28 human sera. Points indicate the mean ± standard error of two technical replicates. (B) Neutralization titers for RSV A2 pseudovirus versus the 28 human sera. RSV pseudovirus neutralization assay titers are the geometric mean of two independent experiments and represent the reciprocal sera dilution at 50% fraction infectivity. RSV A2 full-length virus neutralization assay titers are previously published values and represent the reciprocal sera dilution at 50% fraction infectivity^58^. A correlation plot of these neutralization titers is shown in Figure 3A.

**Supplemental Figure 5.**
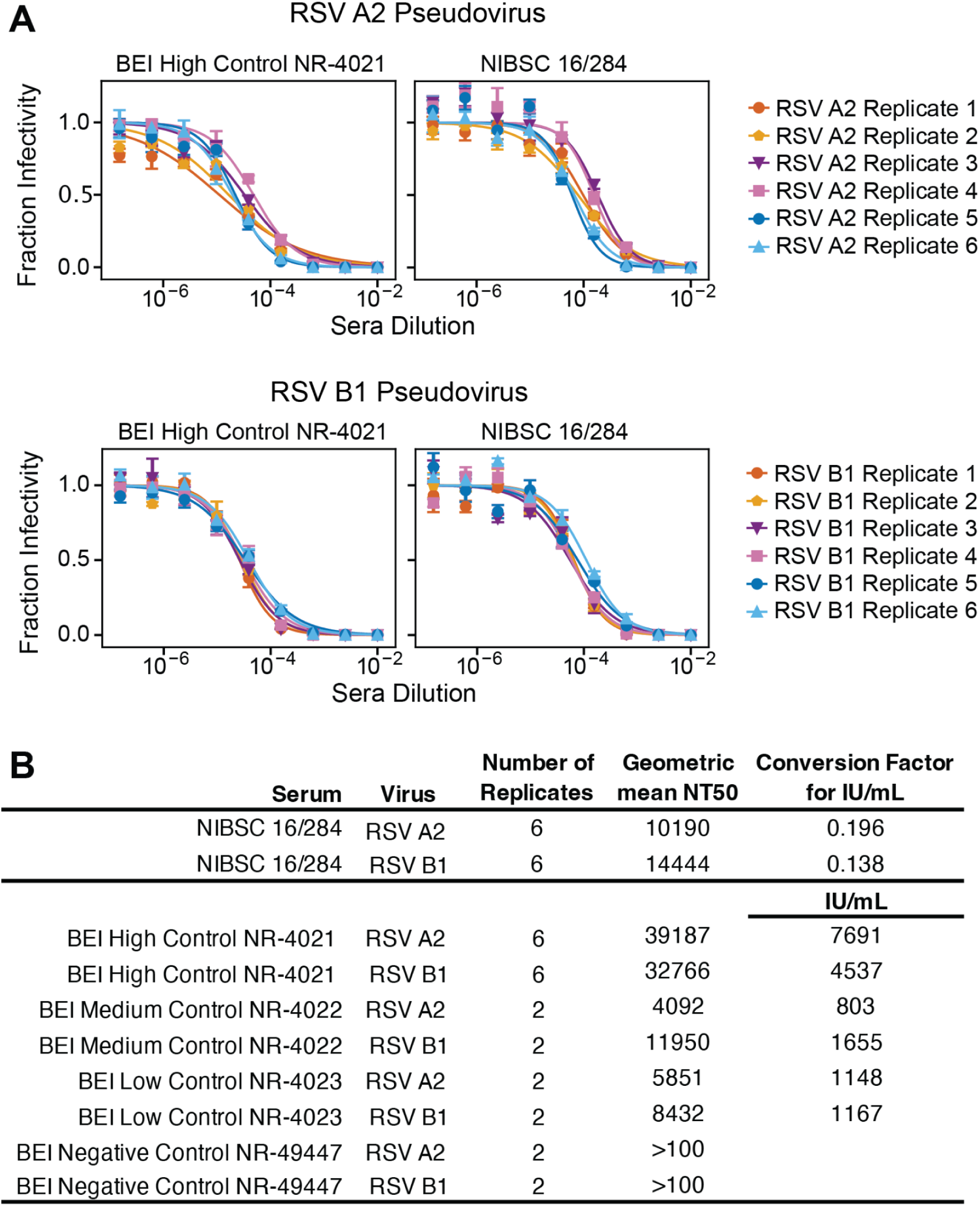
Neutralization measurements are highly reproducible and can be converted to International Units per milliliter. (A) Neutralization curves of pseudoviruses with F and G from the RSV A2 or B1 strain by the BEI High reference serum NR4021 and WHO International Reference Antiserum to RSV (NIBSC 16/284) for six independent experimental replicates performed on three separate days by two different people (Figure 3B shows just two of the replicates for BEI High reference serum NR4021). Points for each replicate indicate the mean ± standard error of two technical repeats of that experiment. (B) Table detailing neutralization titers for the indicated sera and pseudoviruses. The number of replicates indicates the number of independent experiments for which neutralization titers were measured. The geometric mean NT50 represents the reciprocal sera dilution at 50% fraction infectivity. The measured geometric mean titers for the WHO International Reference Antiserum to RSV (NIBSC 16/284) against RSV A2 and B1 were used to calculate a conversion factor based on the standardized potency of these sera (2000/geometric mean titer). This factor converts measurements from our assay into International Units per milliliter (IU/mL)^59,60^. Namely, to convert titers from our RSV pseudovirus assay into IU/ml, those titers should be multiplied by 0.196 for A2, and 0.183 for B1. Neutralization titers for the BEI Resources reference sera were converted to IU/mL by multiplying the measured geometric mean titer by the corresponding conversion factor. Those titers in IU/ml are then plotted versus the known values for the reference sera in Figure 3C.

**Supplemental Figure 6.**
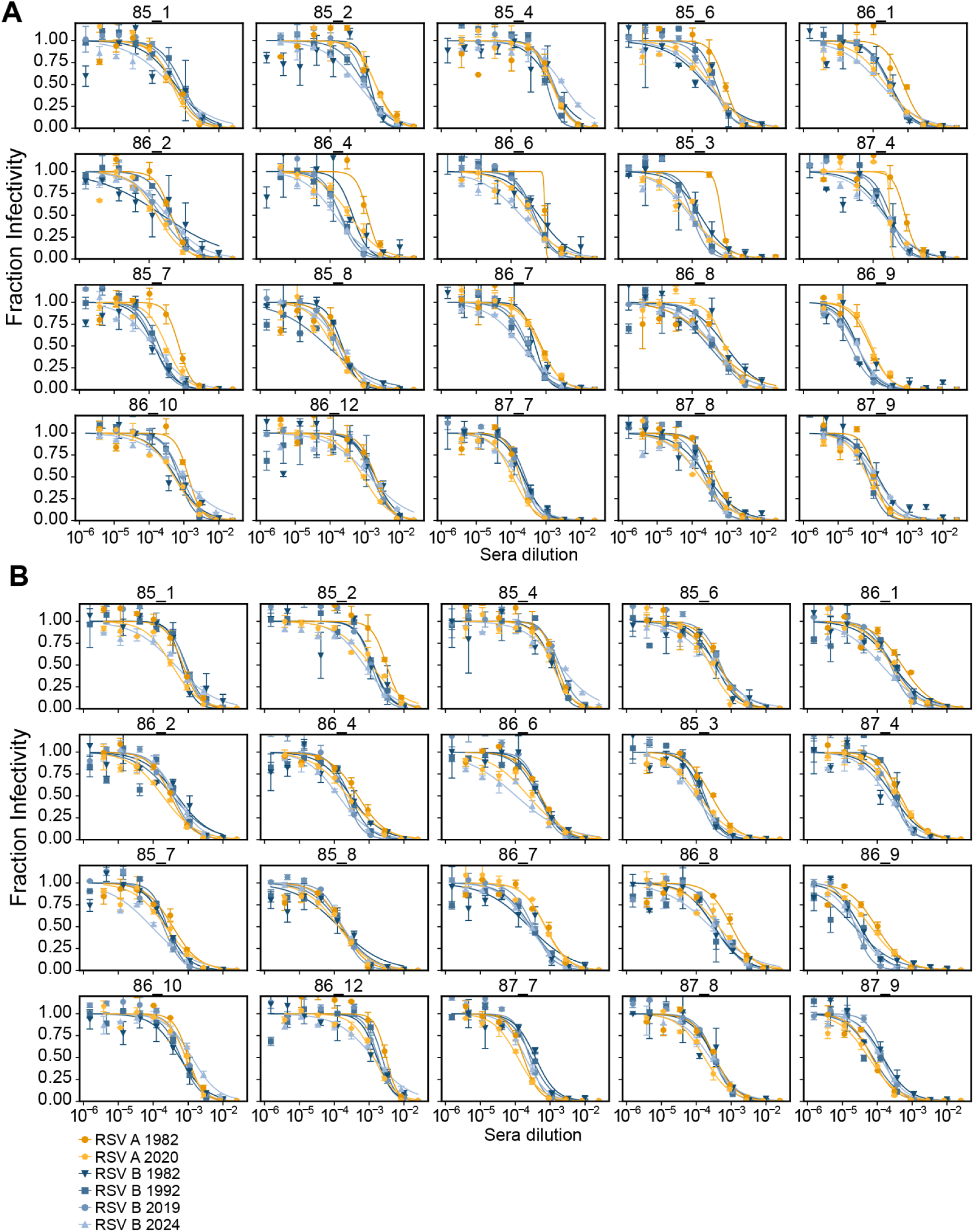
Neutralization curves for historical sera versus RSV pseudoviruses. Neutralization curves for historical serum specimens collected from healthy adults in 1985-1987 (depleted for G-binding antibodies) versus RSV pseudoviruses with G from the Long strain and F from either an historical (eg, 1982) and a recent (eg, 2020-2024) subtype A or B strains. Points indicate the mean ± standard error of technical repeats of experiments performed on the same day. Independent experimental replicates performed on different days are indicated by A and B. A summary plot of the neutralization titers is shown in figure 6B.

**Supplemental Figure 7.**
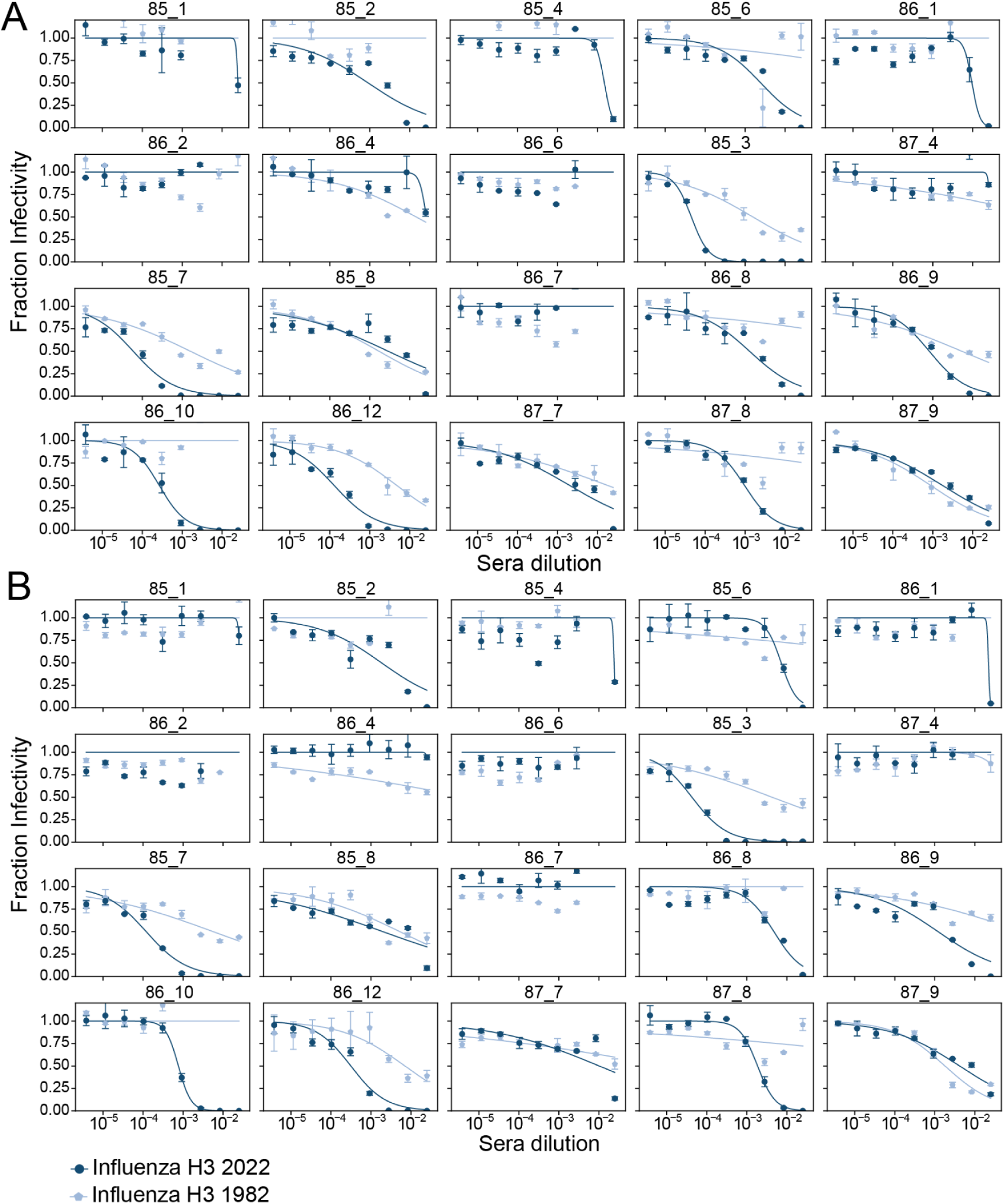
Neutralization curves for historical sera versus pseudoviruses expressing influenza hemagglutinin. Neutralization curves for historical serum specimens collected from healthy adults in 1985-1987 versus pseudovirus expressing hemagglutinin (A/Netherlands/233/1982) and neuraminidase (A/HongKong/1/1968) from historical strains versus pseudovirus expressing hemagglutinin and neuraminidase from a recent (A/Massachusetts/18/2022) strain of human H3N2 influenza. Points indicate the mean ± standard error of technical repeats of experiments performed on the same day. Independent experimental replicates performed on different days are indicated by A and B. A summary plot of the neutralization titers is shown in Figure 6C.

**Supplemental Figure 8.**
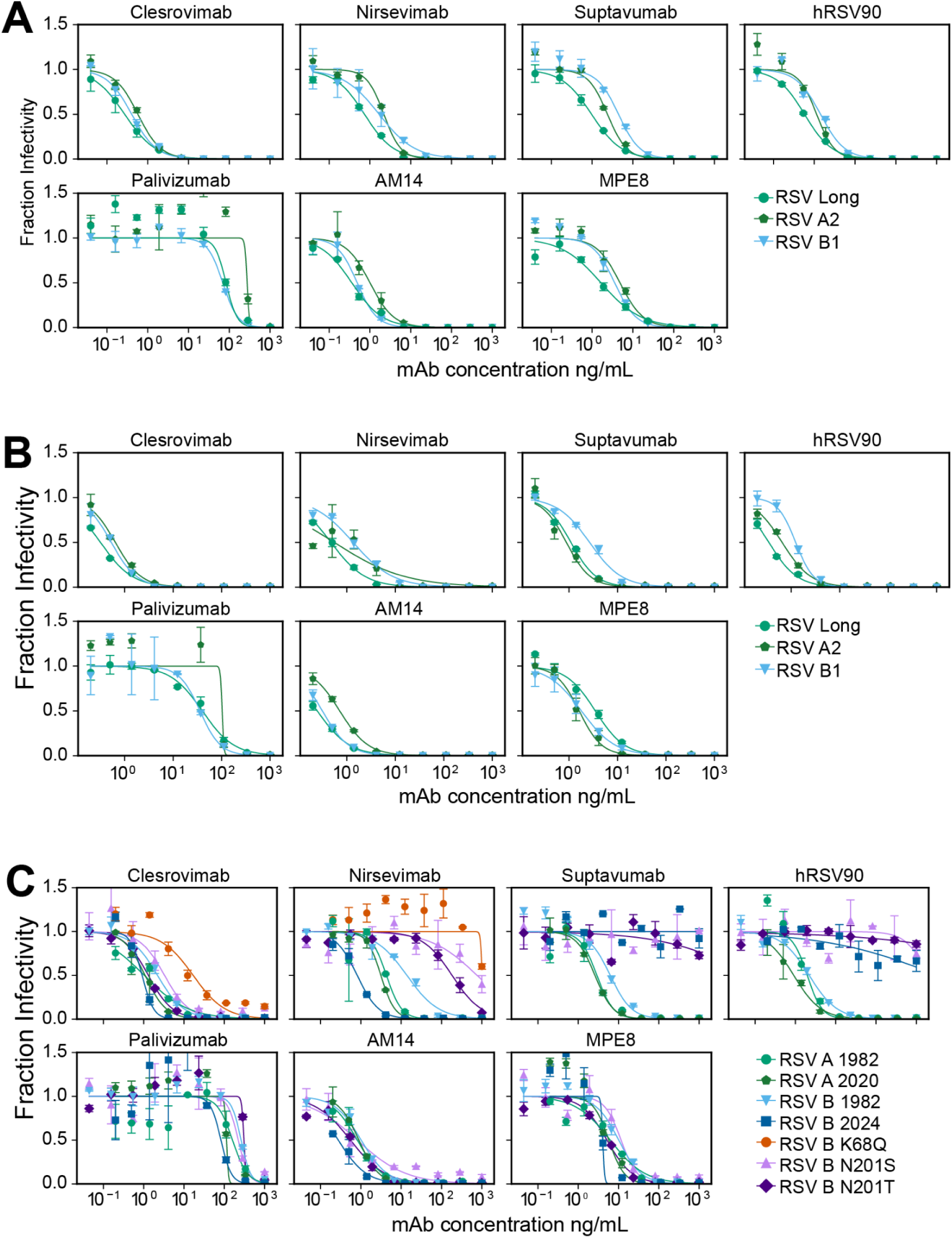
Neutralization curves for monoclonal antibodies versus RSV pseudoviruses expressing F and G from lab-adapted strains. (A and B) Neutralization curves for monoclonal antibodies versus RSV pseudoviruses F and G from lab-adapted strains Long, A2 and B1. Points indicate the mean ± standard error of technical repeats of experiments performed on the same day. A and B represent independent experimental replicates performed on different days. (C) Neutralization curves for monoclonal antibodies versus pseudoviruses expressing the indicated F protein paired with Long G. Points indicate the mean ± standard error of technical repeats of experiments performed on the same day. Neutralization curves for these monoclonal antibodies from a separate experiment performed on a different day are in Figure 7.

**Supplemental Figure 9.**
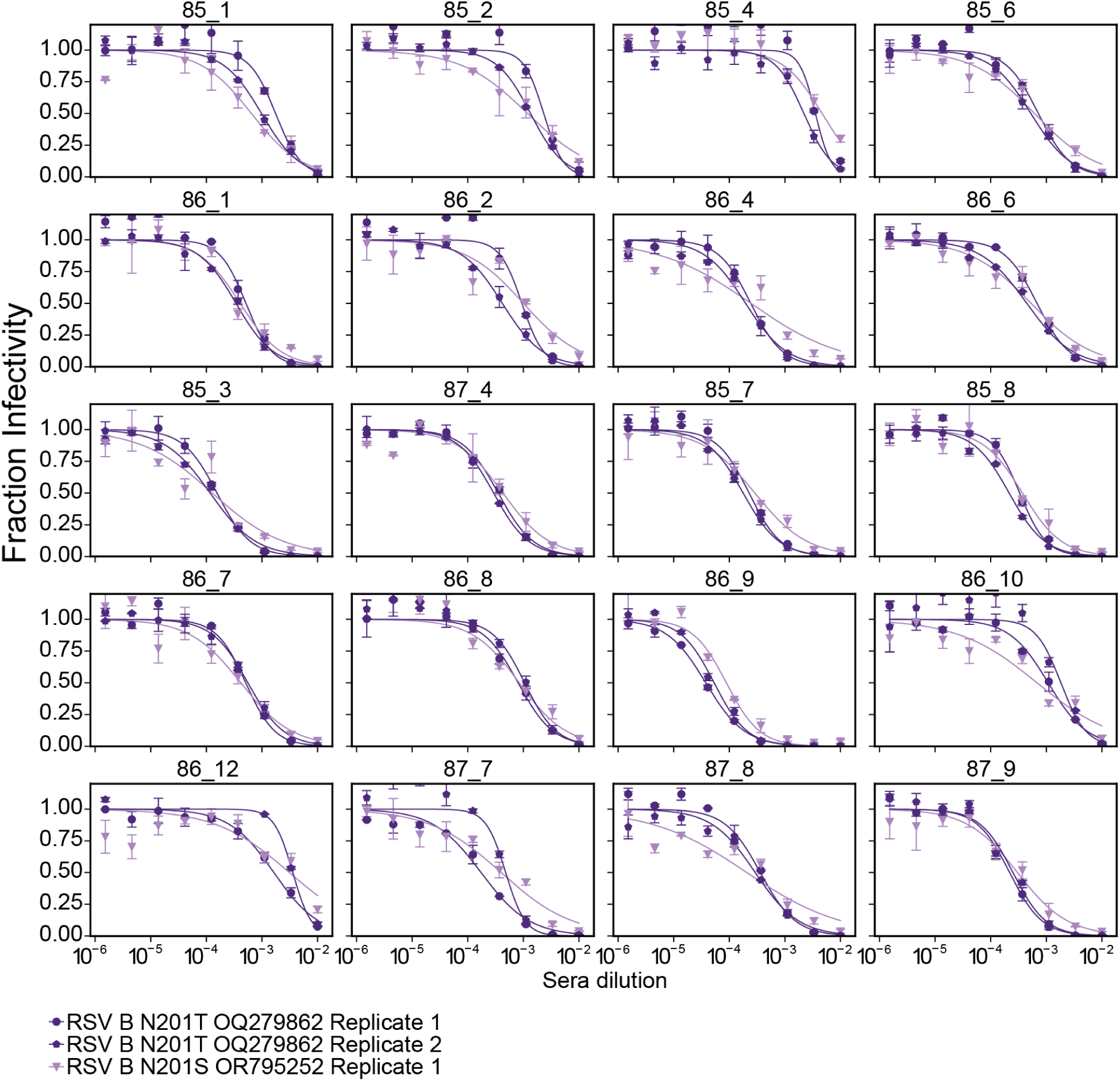
Neutralization curves for historical sera versus RSV pseudoviruses expressing F with nirsevimab resistance mutations. Neutralization curves for historical serum specimens collected from healthy adults in 1985-1987 (depleted for G-binding antibodies) versus RSV pseudoviruses with G from the Long strain and F with known nirsevimab escape mutations. Points indicate the mean ± standard error of technical repeats of experiments performed on the same day. Two independent experimental replicates performed on different days are shown for RSV B N201T OQ279862. One experimental replicate was performed and is shown for RSV B N201S OR795252. A summary plot of the neutralization titers is shown in Figure 7C.

## Supplemental Tables

**Supplemental Table 1.** Sequence accession numbers and pseudovirus names. Plasmid sequences can be found at https://github.com/jbloomlab/RSV-evolution-neut/tree/main/04_plasmid_maps.

| Pseudovirus name | Viral protein | GenBank<br>Accession/GISAID ID | Comments |
| --- | --- | --- | --- |
| RSV Long | RSV Long F | AY911262.1 |  |
|  | RSV Long G | AY911262.1 |  |
| RSV A2 | RSV A2 F | QGW56794.1 |  |
|  | RSV A2 G | KT992094.1 |  |
| RSV B1 | RSV B1 F | AF013254 | also referred to as RSV BWV40167-85 |
|  | RSV B1 G | AF013254 | also referred to as RSV BWV40167-85 |
| RSV A 1982 | RSV A 1982 F | KJ723472 | paired with RSV Long G 31AA CTdel to make RSV pseudovirus |
| RSV A 2020 | RSV A 2020 F | PP495954 |  |
| RSV B 1982 | RSV B 1982 F | MG642043 |  |
| RSV B 1992 | RSV B 1992 F | OK649748 |  |
| RSV B 2019 | RSV B 2019 F | PP352433 |  |
| RSV B 2024 | RSV B 2024 F | PP660445 |  |
| RSV B K68Q<br>MN365362 | RSV B K68Q MN365362 F | MN365362 |  |
| RSV B N201S<br>OR795252 | RSV B N201S OR795252 F | OR795252 |  |
| RSV B N201T<br>OQ279862 | RSV B N201T OQ279862 F | OQ279862 |  |
| Influenza H3 2022 | A/Massachusetts/18/2022<br>(H3N2) HA | EPI2413620 | GISAID ID |
|  | A/Massachusetts/18/2022<br>(H3N2) NA | EPI2413618 | GISAID ID |
| Influenza H3 1982 | A/Netherlands/233/1982<br>(H3N2) HA | EPI545279 | GISAID ID |
|  | A/HongKong/1/1968 (H3N2)<br>NA | AF348184.1 | GenBank ID |
| VSV-G | VSV-G |  | addgene plasmid<br>#12259 |

**Supplemental Table 2.** Sera identifiers, collection year and age at collection for the 20 historical sera specimens.

| sera ID | collection year | patient age |
| --- | --- | --- |
| 85_1 | 1985 | 27 |
| 85_2 | 1985 | 20 |
| 85_3 | 1985 | 26 |
| 85_4 | 1985 | 23 |
| 85_6 | 1985 | 40 |
| 85_7 | 1985 | unknown |
| 85_8 | 1985 | unknown |
| 86_1 | 1986 | 50 |
| 86_10 | 1986 | unknown |
| 86_12 | 1986 | unknown |
| 86_2 | 1986 | 23 |
| 86_4 | 1986 | 25 |
| 86_6 | 1986 | 32 |
| 86_7 | 1986 | unknown |
| 86_8 | 1986 | unknown |
| 86_9 | 1986 | unknown |
| 87_4 | 1987 | 22 |
| 87_7 | 1987 | unknown |
| 87_8 | 1987 | unknown |
| 87_9 | 1987 | unknown |

## Methods

### Data availability

We recommend readers visit https://github.com/jbloomlab/RSV-evolution-neut for well-documented links to visualizations and raw data.

- The full computational analysis of the neutralization data is available on GitHub repository https://github.com/jbloomlab/RSV-evolution-neut/tree/main/02_notebooks
- Visualizations https://jbloomlab.github.io/RSV-evolution-neut/ and https://github.com/jbloomlab/RSV-evolution-neut/tree/main/03_output
- The fraction infectivies used to calculate sera neutralization curves available at https://github.com/jbloomlab/RSV-evolution-neut/tree/main/01_data
- Interactive versions of the RSV F protein phylogenetic trees used in this study are at https://nextstrain.org/community/jbloomlab/RSV-evolution-neut@main/RSV-A-F and https://nextstrain.org/community/jbloomlab/RSV-evolution-neut@main/RSV-B-F.

### Ethics statement

The human sera from the 1980s is from the Infectious Disease Sciences Biospecimen Repository at the Vaccine and Infectious Disease Division (VIDD) of the Fred Hutchinson Cancer Center. These sera were collected from prospective bone marrow donors with approval from the Fred Hutch’s Human Subjects Institutional Review Board. We obtained the human sera previously tested for RSV neutralizing titers using a live virus assay^58^ from the University of Washington Laboratory Medicine with human subjects exemption determination by the Fred Hutch’s Human Subjects Institutional Review Board and not human subjects research determination by the University of Washington Human Subjects Division. All sera are fully de-identified.

### Sera samples

The following reagent was obtained through BEI Resources, NIAID, NIH: Panel of Human Antiserum and Immune Globulin to Respiratory Syncytial Virus, NR-32832. The WHO 1st International Standard Antiserum to Respiratory Syncytial Virus 16/284 was obtained from NIBSC, product number 16/284. See Supplemental Table 2 for available metadata on historical sera samples used in this study and the recently published study by Piliper et al.^58^ for sera used for pseudovirus assay validation. Pooled human sera collected in 2021 was purchased from the Program in Immunology at the Fred Hutch Cancer Center. All sera was heat inactivated at 56°C for 1 hr and then stored long term at -80°C. For assays assessing sera neutralization of influenza HA and NA expressing pseudovirus, sera was treated with receptor destroying enzyme as described previously^93^.

### Antibodies

The RSV monoclonal antibodies palivizumab^73^, nirsevimab^22^, suptavumab^30^, clesrovimab^94^ and hRSV90^77^ were produced by Genscript as human IgG1 kappa isotypes. AM14^75^ and MPE8^76^ were kindly provided by Neil King at the University of Washington. The sequences for heavy and light chains are at https://github.com/jbloomlab/RSV-evolution-neut/blob/main/03_output/summary_tables/RSV_mAb_AAseq.csv. Sequences were obtained from the referenced publications, structures in the Protein Data Bank or original patents. hRSV90 amino acid sequence was obtained from the Protein Data Bank (PDB) 5TPN. Palivizumab originated from patent US6955717B2.

### Cell line handling

All cell lines were cultured in D10 media (Dulbecco’s Modified Eagle Medium supplemented with 10% heat-inactivated fetal bovine serum, 2 mM l-glutamine, 100 U/mL penicillin, and 100 mg/mL streptomycin) and cultured at 37°C with 5% CO2.

### Creation and validation of the 293T-TIM1 cells

Human TIM1 (GenBank: AAC39862.1) followed by 2A peptide and mTagBFP2 was cloned into the pHAGE2 lentivirus vector. TIM1 lentivirus pseudotyped with VSV-G was rescued from 293T cells and used to transduce 293T cells. Note that low titers of this virus were obtained probably due to TIM1 binding virions to producer cells, but we did obtain sufficient titers for transduction. After picking and expanding a single 293T clone, immunostaining was performed to confirm the expression of TIM1 (see Supplemental Figure 2A). For this immunostaining, 293T and 293T-TIM1 cells were digested by trypsin and resuspended into single cells with 1% BSA. Cells were stained with anti-TIM1 primary antibody (AF1750-SP, R&D Systems) diluted in 1% BSA at 1:100 for 1 hour on ice. After three washes with PBS, cells were stained with 488-conjugated donkey anti-goat IgG secondary antibody (A-11055, ThermoFisher) diluted in 1% BSA at 1:1000 for 1 hour on ice. After three washes with PBS, cells were resuspended in 1% BSA for flow cytometry analysis.

### Phylogenetic analysis of RSV F

NextStrain RSV phylogenetic trees are maintained and available at nextstrain.org/rsv^95^. These trees display a subsample of available sequences and are updated over time. To visualize all available sequences at the time of our analysis without subsampling, we utilized the workflow at https://github.com/nextstrain/rsv and removed the subsampling steps to construct subtype A and B F trees containing all sequences as of May 9, 2024 (Figure 5). Interactive versions of these trees are available at https://nextstrain.org/community/jbloomlab/RSV-evolution-neut@main/RSV-A-F and https://nextstrain.org/community/jbloomlab/RSV-evolution-neut@main/RSV-B-F.

### Plasmids encoding viral entry proteins

The RSV F and G’s used in experiments as well as the influenza HA and NA’s are detailed in Supplemental Table 1 including accession numbers. For RSV G’s we deleted 31 amino acids from the N-terminal cytoplasmic tail as shown in Figure 1. RSV Long F and G codon optimized sequences were obtained from a previously published study^50,51^. All other RSV F and G sequences were human codon optimized using a tool by GenScript found at https://www.genscript.com/tools/gensmart-codon-optimization. Codon optimized sequences were then modified to remove homopolymers (>5 nucleotides) and premature poly A signals (AATAAA) which has previously been shown to impact RSV F protein synthesis from transfection^96,97^.

The A/Netherlands/233/1982 (H3N2) HA, A/Massachusetts/18/2022 (H3N2) HA, and A/Massachusetts/18/2022 (H3N2) NA sequences were human codon optimized by GenScript. The A/HongKong/1/1968 (H3N2) NA sequence was human codon optimized by Twist. No further modifications were made to these codon optimized sequences.

We then had the genes synthesized commercially and cloned them into an HDM/CMV driven expression plasmid 27_HDM-tat1b after cutting with NotI and HindIII with the NEBuilder Hifi DNA Assembly Kit incubating at 50°C for 30 min to 1 hour, and transformed into Stellar competent cells (Takara, Cat. # 636763) or NEB 5-alpha Competent E. coli (NEB, Cat. # C2987H). All plasmids were confirmed by whole plasmid sequencing by Plasmidsaurus. The resulting plasmids are available linked in Supplemental Table 1 and the full plasmid maps are at https://github.com/jbloomlab/RSV-evolution-neut/tree/main/04_plasmid_maps.

We generated N-terminal cytoplasmic tail deletions to G by PCR amplification from the full length RSV Long G using primers to delete the desired number of amino acids. C-terminal cytoplasmic tail deletions to F were generated by PCR amplification from the full length RSV Long using primers to introduce a stop codon. These gene fragments were cloned into the expression vector as described above.

All plasmid maps are available at https://github.com/jbloomlab/RSV-evolution-neut/tree/main/04_plasmid_maps. The lentivirus helper plasmids are available from AddGene: HDM-tat1b product ID 204154, pRC-CMV-Rev1b product ID 20413, HDM-Hgpm2 product ID 204152^98^. The VSV-G expression plasmid used as a positive control is also available from AddGene: HDM_VSV_G product ID 204156. A lentiviral backbone plasmid that uses a CMV promoter to express luciferase followed by an IRES and ZsGreen is available from BEI Resources (BEI catalog number NR-52516)^52^.

### Generation and titration of pseudotyped lentiviral particles encoding luciferase and ZsGreen

We generated pseudovirus expressing RSV F and G and encoding luciferase and ZsGreen as shown in Figure 1. To make RSV pseudovirus, 293T cells were plated the day before transfection in D10 to achieve ∼70% confluency on the day of transfection. Cells were plated in either 6-well dishes (600,000 cells per well), 10 cm dishes (4,000,000 cells) or 15 cm dishes (11,000,000 cells) depending on the desired volume of viral supernatant. 16-24 hours after plating, 293Ts were transfected using BioT (Bioland Scientific) with the appropriate ug of plasmid DNA designated in the product manual. For example in a single well of a 6-well dish, 2 ug total of DNA was transfected split up between 1 ug of the lentiviral backbone plasmid encoding luciferase and ZsGreen, 200 ng of RSV F expression plasmid, 100 ng RSV G expression plasmid and 233 ng each of lentiviral helper expression plasmids 26_HDM-Hgpm2 (Gagpol), 27_HDM-tat1b (Tat), and 28_pRC-CMV-Rev1b (Rev). If producing in larger plates, DNA was scaled up accordingly, keeping the proportions the same between plasmids. After around 48 hours the supernatant was filtered using a 0.45 um syringe filter. Aliquots of viral stocks were then labeled and flash frozen using a prechilled CoolRack CF45 Cooling Block (Fisher Cat. #UX-04392-51) on dry ice until frozen solid (around 15 minutes) before transferring to -80°C for long term storage.

To titrate these pseudoviruses (Figure 1 and Figure 2), we seeded 60,000 target cells (293T or 293T-TIM1 cells) per well in 100 uL D10 on 96-well plates the day before infection to achieve 90% confluency at infection. We plated in white-walled, clear bottom 96-well plates for luciferase measurements and clear 96-well plates for ZsGreen measurement by flow cytometry. 16-24 hours after plating aliquots of RSV pseudovirus were thawed in a 37°C water bath and serially diluted in 96-well plates. Technical replicates were performed within each experiment. We then added 100 uL of diluted virus per well to the cells for infections. If determining viral titer by flow cytometry, prior to transferring virus dilutions, 2-4 wells of the 96 well plate were trypsinized and counted to determine cell count per well at the time of infection for TU/mL calculations. Titer plates were harvested at 48-72 hours following infection.

If determining titers by luciferase, we measured RLU using the Bright-Glo Luciferase Assay System (Promega, Ref. No. E2620). Briefly, 150 uL of media was aspirated from wells to leave ∼30uL per well before adding 30 uL of BrightGlo reagent per well to each well of the white 96 well plate for a 1:1 volume ratio in each well. To reduce background luminescence, we placed a white sticker to mask the clear bottoms of all wells or transferred to opaque white plates. We measured luminescence activity by plate reader. For readings that were in the linear range, we normalized counts to volume to generate RLU/uL. For each pseudovirus, we calculated an RLU/uL from technical replicates by taking the mean of all individual replicates.

If determining titers using flow cytometry based on percent positive ZsGreen cells, wells with ∼1-10% green cells were estimated by eye, trypsinized from the plate, spun down in a V-bottom 96 well plate and washed 3 times with 1-3% BSA in PBS. Washed cells were then read on a flow cytometer and % ZsGreen positive cells was determined based on gating of uninfected wells in FlowJo. This along with the cell count per well on the day of infection was used to calculate TU/mL (transducing units per mL).

We generated pseudovirus expressing influenza HA and NA using a similar protocol as described above with a few modifications. 293T cells were plated in 10 cm dishes the day before transfection in D10 to achieve ∼70% confluency on the day of transfection. 16-24 hours after plating 293Ts were transfected using BioT (Bioland Scientific) with 11.25 ug of DNA total per dish, 5 ug of lentiviral backbone encoding luciferase and ZsGreen, 1.25 ug each of lentiviral helper plasmids, 1.25 ug HA, 0.25 ug NA and 1 ug of the HAT protease that activates HA. 12 to 16 hours post transfection media was swapped from D10 to IGM (influenza growth media)^93^. Viral supernatant was collected around 48 hours post transfection by syringe filtering through a 0.45 um filter. Aliquots were stored at -80°C. To determine viral titers by luciferase, MDCK-SIAT1 cells^99^ were plated on the day of infection around 2 hours before infection by adding 15,000 cells per well in 50 uL of NAM (neutralization assay media, consisting of Medium-199 supplemented with 0.01% heat-inactivated FBS, 0.3% BSA, 100 U/mL penicillin, 100 ug/mL streptomycin, 100 ug/mL calcium chloride, and 25 mM HEPES)^100^ supplemented with 2.5 ug/mL Amphotericin B, which increases pseudovirus titers^98^. Serial dilutions of pseudovirus were then set up in NAM supplemented with 500 nM of oseltamivir, which inhibits neuraminidase from cleaving sialic acids on target cells, therefore making entry more efficient. The dilution plate was incubated on ice for 20 min for oseltamivir to bind. Then 100 uL of each serial dilution was added to previously plated cells. After adding pseudovirus, the MDCK-SIAT1 plate was incubated for 48 hours. To read titer for luciferase activity BrightGlo was added and plates were read as described above for RSV.

### RSV F- and G-binding antibody depletion from sera and validation of depletion

We depleted sera of F- or G-binding antibodies as shown in Figure 4. We obtained commercially available His-tagged, soluble RSV G protein based on the Long strain from SinoBiological (Cat: 40041-V08H). Soluble RSV F protein based on the Long strain both unmodified and stabilized in the pre-fusion conformation using the SC-DM mutations as previously described^101^ were produced by removal of the transmembrane and cytoplasmic tail domains, and addition of a trimerization domain^102^ and N-terminal His-tag. The proteins were produced with pCMV/R vectors using endotoxin-free DNA in ExpiHEK293F cells grown in suspension using Expi293F expression medium (Life Technologies) at 33°C, 70% humidity, 8% CO_2_ rotating at 150 rpm. The cultures were transfected using PEI-MAX (Polyscience) with cells grown to a density of 3.0 million cells per mL and cultivated for 3 days. Supernatants were clarified by centrifugation (5 minutes at 4000 rcf), addition of PDADMAC solution to a final concentration of 0.0375% (Sigma Aldrich, #409014), and a second centrifugation (5 minutes at 4000 rcf). The secreted proteins were purified from cell supernatant by IMAC, followed by SEC on a Superdex 200 10/300 GL (GE Healthcare) equilibrated in 50 mM Tris pH 7.4, 250 mM NaCl, 50 mM Glycine, and 5% v/v Glycerol.

Magnetic Dynabeads for His-tag isolation and pulldown (Invitrogen Cat. # 10104D) were aliquoted (100 uL per condition) and washed once with PBS in a magnetic stand. After removal of supernatant washed beads were resuspended in 10 ug protein (or PBS for mock conditions). Beads were incubated with protein for 10 minutes at 4°C on a rocker to allow his tagged protein to bind with the magnetic beads. After incubation the beads were washed twice and supernatant was discarded. Beads were then resuspended in 100 uL sera. Sera and bead mixture was incubated on a rocker at 4°C overnight. After overnight incubation, samples were placed on a magnetic rack for 2 minutes to separate antibodies bound to protein conjugated beads from supernatant. The supernatant was collected and a second round of depletion was performed. Depleted sera was stored at -80°C.

We validated depletion of RSV F- or G-binding antibodies from sera by ELISA (Figure 4B). Immunlon 2HB 96-well plates (Thermo Scientific 3455) were coated with 100 uL of RSV Long F, preF or G protein at 1ug/mL for 30 minutes at 37°C in PBS. Unbound protein was then removed by washing three times in PBS + 0.1% Tween-20 (PBS-T). Plates were blocked for one hour at room temperature in 200 uL per well 5% (w/v) dried milk in PBS-T. Blocking buffer was removed and sera/mAbs from serially diluted sera was added (100 uL per well) and incubated at 37°C for one hour. After incubation plates were washed three times in PBS-T and then incubated for one hour at 37°C in secondary antibody HRP-conjugated goat anti-human IgG (Bethyl Labs, Cat. #A80-104P) at a dilution of 1:2000 in blocking buffer (50 uL per well). Secondary antibody was washed off with three washes in PBS-T. Plates were developed with 100 uL TMBE for four minutes and stopped with 100 uL 1N HCL before reading absorbance (450 nm) on a plate reader.

### Pseudovirus neutralization assays

Neutralization assays with pseudovirus were performed as previously described with the modifications described below^52,70^. To perform neutralization assays using pseudoviruses expressing RSV F and G, we plated 293T-TIM1 cells (60,000 per well) in 96-well white-walled clear-bottom tissue culture treated plates 16-24 hours prior to setting up the neutralization assay. Serum samples were diluted 1:10 or 1:50 in D10 and then serially diluted 3-fold for a total of 9 dilutions per serum. Monoclonal antibodies were diluted to 1000 ng/mL with 9 total 3 or 3.5-fold serial dilutions. 50 uL of these diluted serum or mAb samples were then added 96-well ‘set-up’ plates in duplicate (2 columns) per serum sample. Pseudoviruses expressing RSV F and G were then diluted the appropriate amount to achieve 600,000 to 2,000,000 RLU per well, these dilution factors varied per virus due to differences in titer (Figure 1 A,B) but were always diluted at least 2 fold (Supplemental figure 3). 50 uL of diluted virus was then added to assay ‘set-up’ plates that contain 50 uL of serum or mAb (note that pseudovirus is therefore diluted at least 4-fold after this step, Supplemental Figure 3) as well as 50 uL of virus only added to 50 uL of D10 as a virus only control and ‘set-up’ plates were incubated for 1 hour at 37°C with 5% CO2. After the incubation 100 uL per well of the set up plate was transferred to the plated target cells. These plates were then incubated for 48 hours before reading the luciferase signal as described above for pseudovirus titration.

Neutralization assays for pseudoviruses expressing influenza HA and NA were performed similarly to those using RSV pseudoviruses with a few differences. As described for titration of viruses, MDCK-SIAT1 cells were plated in clear-bottom white-walled 96-well tissue culture plates 2 hours before infection. Cells were plated in 50 uL of NAM supplemented with amphotericin B. Serum samples were first treated with RDE which results in a 1:4 dilution of sera, this initial dilution was accounted for when setting up serially-diluted sera with starting dilution of 1:10. Serum dilutions and virus dilutions were done in NAM and virus dilutions were supplemented with 500 nM of oseltamivir. The rest of the assay was performed as described above for RSV pseudotyped lentivirus.

Fraction infectivity was calculated as compared to a no-serum or no-antibody well. Neutralization curves were then fit to the data using the neutcurve package https://jbloomlab.github.io/neutcurve^103^.

